# The Type VI Secretion System Antifungal Effector Tfe2 Inhibits Protein Translation and Drives Hyperactivation of TORC1

**DOI:** 10.1101/2025.11.20.689513

**Authors:** Katharina Trunk, Genady Pankov, Alex Anderton, Viktoras Stony, Andrew Frey, Jennifer Haggarty, Ian Leaves, Pantelitsa Papakyriacou, Mengchun Li, Vivek Manthri, Flávia Zimbres, Paul W Denny, Phillip D Whitfield, Christian Hacker, Neil AR Gow, Colin Rickman, Christopher M Grant, Matthias Trost, Sarah J Coulthurst, Janet Quinn

## Abstract

Type VI Secretion Systems (T6SS) are utilised by many bacteria to deliver toxic effectors into neighbouring bacterial, fungal or host cells. Whilst many antibacterial effectors are well characterised, much less is known regarding the identity or mode-of-action of antifungal effectors. Here we combine structural modelling with proteomics and in vivo approaches, to show that the *Serratia marcescens* antifungal effector Tfe2 adopts a novel fold and functions as a potent inhibitor of protein translation leading to hyperactivation of the TORC1 kinase. We show that Tfe2 expression in *Saccharomyces cerevisiae*, or treatment with the protein translation inhibitor cycloheximide, drive identical increases in free intracellular amino acids and hyperactivation of TORC1. This, in turn, triggers the Tfe2 and cycloheximide-mediated rapid turnover of amino acid transporters through stimulating substrate-independent endocytosis. Polysome profiling, however, revealed differences in Tfe2 and cycloheximide-mediated protein translation inhibition, with Tfe2 inhibiting initiation of translation. Tfe2-mediated hyperactivation of TORC1 may also underpin adaptive responses to this effector which include significant remodelling of the lipidome and notable alterations in organelle and cell wall structures. Collectively this study has provided new insight into the mode-of-action of a structurally novel antifungal effector Tfe2.

## INTRODUCTION

Fungi and bacteria are rarely found in isolation and instead exist in polymicrobial communities where multiple synergistic and antagonistic interactions determine community composition and dynamics^1^. Such cross-kingdom interactions are emerging as key determinants in human microbiota composition and health^2, 3^, with a growing number of examples of medically relevant bacterial-fungal interactions^4^. Antagonistic bacterial-fungal interactions have long been recognised ^1^, with those relevant to human health including bacterial manipulation of the shared environment^5^, or through the secretion of diffusible molecules that display antifungal activity or inhibit key fungal virulence traits^6, 7, 8^. In addition, we reported that the opportunistic pathogen *Serratia marcescens* can deploy its Type VI Secretion System (T6SS) as a potent antifungal weapon through injecting anti-fungal toxins directly into fungal competitors. Two dedicated T6SS-elicited antifungal effectors, Tfe1 and Tfe2, were identified that mediate *S. marcescens* killing of model and pathogenic yeasts^9^. This discovery, that a bacterium can employ their T6SS against fungal competitors, revealed a previously uncharacterised mechanism underpinning antagonistic bacterial-fungal interactions.

The T6SS is a large multiprotein structure, widely distributed in Gram-negative bacteria, that functions as a contractile nanomachine to fire toxic effector proteins directly into neighbouring cells. Within the T6SS, a cytoplasmic tubular sheath structure, anchored in the inner and outer bacterial membranes, contracts and expels a needle like structure, decorated with multiple effector proteins, out of the bacterial cell into the target cell^10, 11^. Following breach of the target cell, effectors are released where they induce toxicity by distinct mechanisms. In some cases, the T6SS is used to target higher eukaryotic cells through the secretion of anti-host effectors^12^. However, most characterised T6SS-elicited effectors are antibacterial effectors and include peptidoglycan hydrolases, phospholipases, NADases, ADP-ribosyltransferases, DNA nucleases and deaminases, and pore forming toxins ^13^. Following the discovery that the antibacterial *S. marcescens* T6SS is also deployed against fungal competitors, T6SS-dependent antifungal activity has been identified in a number of environmental^14, 15^ and plant and human pathogenic bacterial species^16, 17, 18, 19^. Thus, T6SS mediated antifungal activity appears to be widespread, targeting a broad range of fungal species from yeasts to filamentous fungi. Despite this, only a few T6SS-elicited antifungal effectors have been identified in addition to *S. marcescens* Tfe1 and Tfe2, which include *Klebsiella pneumonia* VgrG4, *Acinetobacter baumannii* TafE and *Acidovorax citrulli* TseN^16, 20, 21^. Whilst TafE and TseN display DNase activity^16, 21^, and VgrG4 induces mitochondrial fragmentation^20^, the precise mode of action of Tfe1 and Tfe2 remain unclear^9, 22^.

The *S. marcescens* effectors Tfe1 and Tfe2 appear to act via distinct mechanisms against a number of yeast species to cause cell death, with Tfe2 being responsible for the majority of antifungal activity against *Saccharomyces cerevisiae* and *Candida glabrata*, and Tfe1 and Tfe2 both contributing to the killing of *Candida albicans*^9^. These effectors are small proteins that are conserved across a number of bacterial species^22^, however, bioinformatic analysis failed to reveal any conserved domains or predicted functions. Tfe1 was found to drive depolarisation of the fungal plasma membrane independent of pore formation^9^, but the precise mechanism underpinning this remains unclear. Regarding Tfe2, ‘in competition’ proteomics revealed that Tfe2 intoxication triggered the downregulation of several amino acid transporters and proteins in the sulphate assimilation pathway, which culminated in nutrient imbalance and the induction of autophagy^9^. However, the mechanisms underpinning these responses, or whether such responses are direct or indirect responses to Tfe2 toxicity, remain unknown. Thus, there are significant knowledge gaps in our understanding of T6SS-mediated antifungal activity both regarding the identification of effectors and in defining their antifungal mode-of-action.

Here we present a comprehensive analysis of the *S. marcescens* antifungal effector Tfe2. We show that Tfe2-mediated downregulation of amino acid transporters is a prominent early cellular response of *S. cerevisiae* cells to this effector, and that this is a consequence of Tfe2 functioning as a potent inhibitor of protein translation initiation. We demonstrate that Tfe2-mediated translation inhibition drives a rapid increase in free intracellular amino acids, and TORC1 hyperactivation, which likely underpins the rapid substrate-independent endocytosis of amino acid transporters. TORC1 hyperactivation may also contribute to secondary cellular responses to Tfe2-intoxication, including significant changes to the lipidome, organelle structure and fungal cell wall of *S. cerevisiae* cells. The precise mechanism underpinning Tfe2 inhibition of translation remains to be deciphered but bioinformatics, structural modelling and mutagenesis analysis indicate that Tfe2 adopts a novel fold that is structurally conserved across a previously uncharacterised effector family.

## RESULTS

### Structural modelling and site directed mutagenesis indicate that Tfe2 adopts a novel structure and functions via an acid-base hydrolysis mechanism

Despite extensive efforts, soluble recombinant Tfe2 could not be purified preventing structural determination, and thus an AlphaFold model of Tfe2 was generated. This displayed high per-residue confidence across the chain (mean pLDDT 94.5) with only terminal loops showing modestly lower values (Fig. 1a). This model revealed a compact, predominantly α-helical fold, with helices arranged in a twisted bundle stabilised by loop regions and anchored by a short antiparallel β-sheet at the base of a pronounced surface cleft. Electrostatic surface mapping highlights a shallow, negatively charged cavity on one face of the protein, suggestive of a functional site (Fig. 1b).

**Figure 1.**
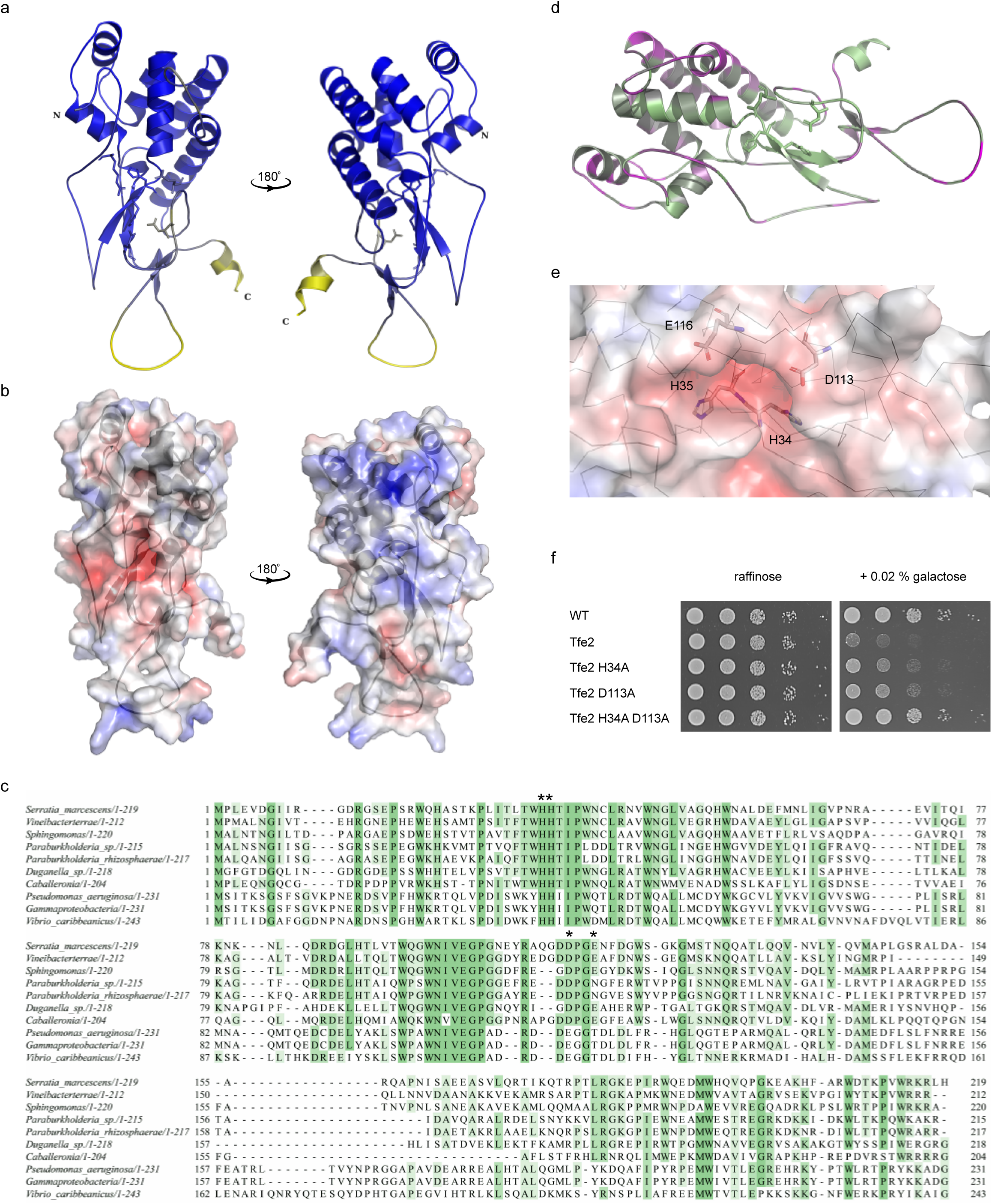
Tfe2 adopts a novel structural fold, representing a previously uncharacterised effector family that likely functions via an acid base hydrolysis mechanism. **(a)** Cartoon representation of the AlphaFold-predicted structure of *S. marcescens* Tfe2 shown in two orientations rotated 180 °C, and coloured in the per-residue pLDDT confidence score, from high confidence (blue) to low (yellow). **(b)** Electrostatic surface representation of Tfe2 (same orientation as in panel a), coloured by charge distribution, from negative (red) to positive (blue). **(c)** Multiple sequence alignment of Tfe2 and bacterial homologues, coloured by percentage identity (green = conserved). Conservation of predicted key catalytic residues is indicated with asterisks. **(d)** Ribbon representation of Tfe2 coloured by evolutionary conservation as calculated by ConSurf. The structure is coloured from most conserved residues (green) to least conserved (magenta). **(e)** Close-up view of the negatively charged surface cleft, showing the side chains of conserved polar and charged residues (His34, His35, Asp113, Glu116) as sticks arranged in geometry consistent with acid-base hydrolysis. **(f)** Growth of *S. cerevisiae* K699 chromosomal integration strains carrying the empty promoter construct (WT) or constructs directing the expression of the wild type Tfe2, single amino acid variants (Tfe2 H34A, Tfe2 D113A) or the double mutant variant (Tfe2 H34A D113A) on media containing raffinose or galactose for repression or induction, respectively, of gene expression.

Sequence alignment of *Sm*Tfe2 homologues (Fig. 1c) and conservation mapped onto the structure (Fig. 1d) show that residues lining the cavity are strongly conserved. This cavity contains polar and charged residues arranged in a geometry compatible with catalysis (Fig 1e). In particular, the proximity of His34 and His35 to Asp113 and Glu116 is consistent with a classical acid–base hydrolysis mechanism, where histidines may activate a water molecule and acidic residues stabilise the transition state or act as proton donors. Such configuration is characteristic of catalytic dyads present in various hydrolases^23, 24, 25^.

Although motif-based searches using ScanProsite^26^ and structure-based comparisons using the DALI server^27^ failed to identify structurally similar proteins with known function, the overall architecture and surface electrostatics (Fig. 1a, b) are reminiscent of active centres in known hydrolases or oxidoreductases. Similarly, HHpred analysis^28^ did not detect any significant sequence homology to annotated functional domains, supporting the view that Tfe2 represents a structurally conserved but functionally uncharacterised effector family. The predicted negatively charged cavity appears solvent accessible and electrostatically suited to accommodate a positively charged or polar substrate. Taken together, these features suggest that Tfe2 contains a catalytically competent site defined by evolutionarily conserved residues.

To probe the functional relevance of these putative catalytic residues, site-directed mutagenesis was performed. Single alanine substitutions at His34 and Asp113 caused mild attenuation of fungal growth inhibition when conditionally expressed in *S. cerevisiae*, whereas the H34A/D113A double mutant completely abolished Tfe2 activity, indicating a cooperative role in catalysis (Fig. 1f). Immunoblotting of GFP-tagged variants confirmed that loss of activity was not due to reduced protein abundance (Fig. S1), and AlphaFold predictions of mutant structures did not suggest folding issues, supporting a direct catalytic role for these residues.

The conservation of His34, His35, and Asp113 across homologues, along with structurally preserved positioning within the predicted active site, supports a conserved mechanism of action. Glu116, spatially proximate to His35, is also conserved in several homologues and may contribute to catalysis or transition state stabilisation.

In conclusion, the predicted structure of Tfe2 adopts a novel fold that is structurally conserved across a previously uncharacterised effector family. In addition, taken together, the structural modelling, mutagenesis data, and conservation across homologues support a model in which Tfe2 functions via an acid-base hydrolysis mechanism mediated by a His-Asp dyad, with auxiliary contributions from neighbouring residues such as His35 and Glu116.

### Proteomics analysis identifies early and adaptive responses of yeast cells to Tfe2 intoxication

As challenges in protein purification of Tfe2 did not allow for enzyme-based discovery methods, we performed a detailed analysis of proteomic changes upon conditional expression of Tfe2 in the model yeast *S. cerevisiae*.

Previous ‘in competition’ proteomics analysis following co-culture of *C. albicans* with *S. marcescens* wild-type and effector mutants, revealed Tfe2 to cause a downregulation of amino acid transporters, and enzymes within the sulphate assimilation pathway, leading to the induction of nitrogen starvation and autophagy ^9^. Whilst this analysis allowed for identification of Tfe2 induced proteomic changes in a physiological relevant setting, we speculated that affected cells would be subject to fluctuation in cellular effector concentration with differing intoxication times, which would present as a mix of primary and secondary responses. To differentiate between initial and adaptive *S. cerevisiae* responses to Tfe2 we used the galactose-inducible expression system and analysed its proteome at onset (0 h), short term (1 h) and medium term (6 h) induction, comparing to the WT strain grown in the absence and presence of galactose.

Principal component (PC) and volcano plot analysis indicated good overlap of both WT and Tfe2-expressing strains at onset of induction, with their proteomes moving out and populating distinct areas on the PC plot over time (Fig. 2a, b). Initial proteomics analysis was carried out using Limma in R and the resulting dataset (Table S1) was transferred to Proteus for ANOVA analysis. Hierarchical clustering of the 2007 ANOVA positive protein responses into 16 protein clusters (Fig. 2c), identified 11 clusters with significantly altered protein profiles, of which five clusters displayed WT specific responses to the addition of the preferred carbon source galactose. These represent proteins involved in energy metabolism, including downregulation of proteins of the fatty acid and amino acid metabolic processes and upregulation of proteins involved in carbohydrate metabolic processes (clusters 6, 7, 14, 9 and 12, Fig. 2d, Table S2). These changes were not elicited in the Tfe2-expressing cells, in accordance with its previously described metabolic and growth inhibitory activity^9^. The remaining six clusters featured proteins specifically downregulated upon Tfe2 intoxication and can be separated into early (clusters 3 and 8) and adaptive responses (clusters 2, 4, 10 and 11, Fig. 2d, Table S2), including nutrient transport and ubiquitin-dependent transporter endocytosis, ribosomal subunits and translation related processes, sterol and sphingolipid metabolic processes and cell wall-related processes. The clear distinction between growth-related responses of the WT to the inducing agent galactose, which were absent in the Tfe2-intoxicated cells, and the cellular responses to the toxin, prompted us to focus our further analysis on the differential proteome of Tfe2 cells under non-inducing and inducing conditions.

**Figure 2.**
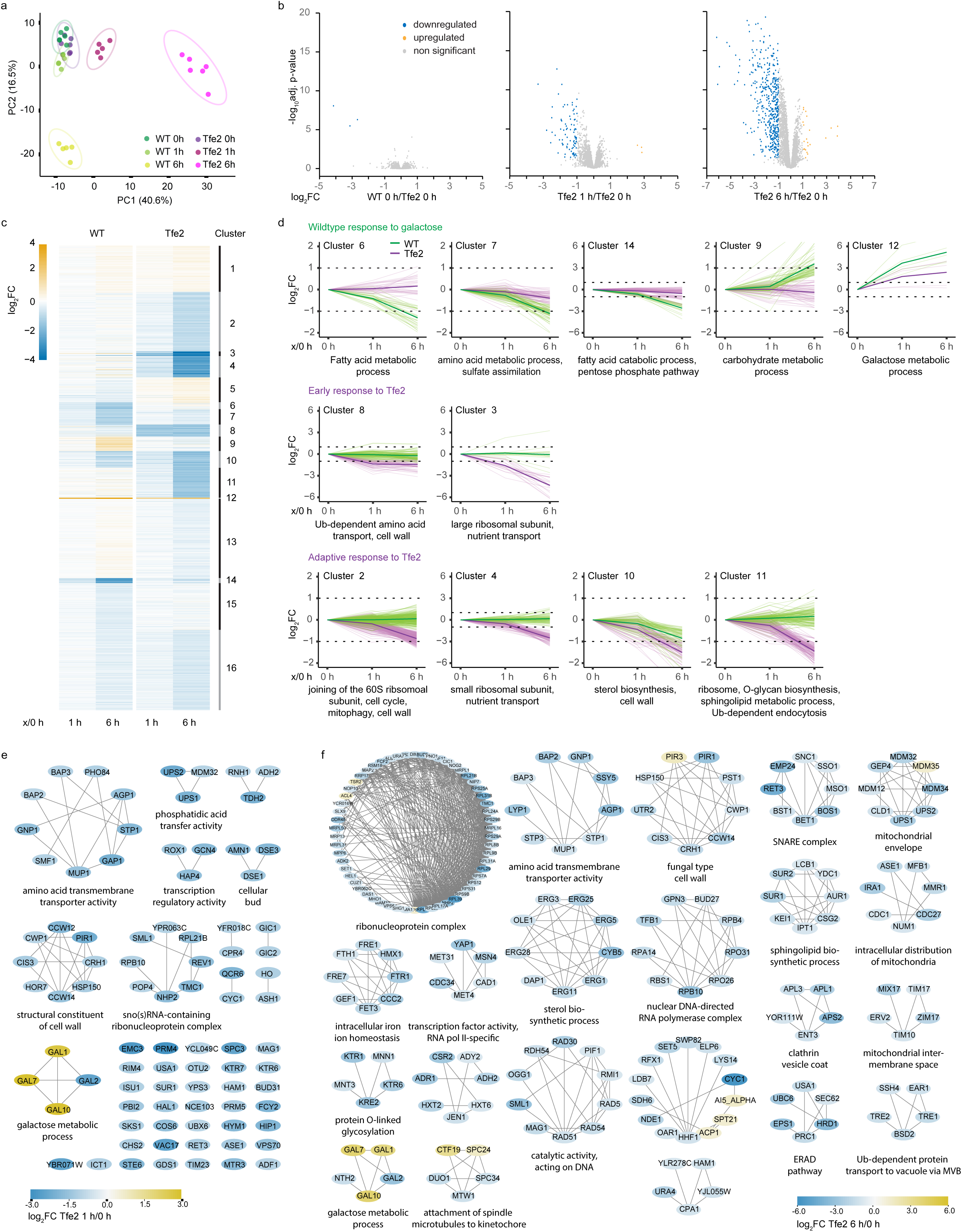
Proteomics analysis of the early and adaptive yeast response to Tfe2 intoxication. **(a)** Principal component (PC) analysis of the proteomes of *S. cerevisiae* control (WT, green) and Tfe2-expressing (purple) strains before (0 h) and after 1 and 6 h of induction. **(b)** Volcano Plots showing differential expression of proteins expressed in the *S. cerevisiae* control strain (WT) relative to the Tfe2-harbouring strain at 0 h, and after 1 h and 6 h of Tfe2-expression, as compared to the un-induced (0 h) Tfe2 strain. Statistically significant downregulated proteins are in blue, upregulated proteins in yellow and unregulated proteins in grey (log_2_ fold change >1, p-values adjusted for false discovery rate for multiple comparisons using the Benjamini-Hochberg test, p <0.05) **(c)** Hierarchical clustering of the 2007 ANOVA positive protein responses (WT 1h and 6h/WT 0h, Tfe2 1h and 6h/Tfe2 0h; p-values adjusted for false discovery rate using a multiple comparison Benjamini-Hochberg test, p <0.0001) into 16 protein clusters (unsupervised K means clustering). Differential protein expression is coloured from downregulated (log_2_FC = −4, blue) to upregulated proteins (log_2_FC = 4, yellow). **(d)** Single clusters of the hierarchical clustering analysis in c), depicting differential protein expression of WT (green) and Tfe2-expressing (magenta) strains, with a mean WT or Tfe2 response at 1 or 6 h >2-fold. Clusters are grouped into WT response to galactose (upper panel), early response to Tfe2 (middle panel) and adaptive response to Tfe2 (lower panel). **(e,f)** Functional enrichment analysis by topological clustering based on interactions of differentially regulated proteins after 1h (e) and 6h (f) Tfe2-intoxication as compared to uninduced conditions using a Markov Cluster algorithm (MCL = 3). Shown are all clusters retrieved (e) and clusters with a minimum of 5 interacting proteins, respectively (f), with differential protein expression being coloured from downregulated (blue) to upregulated proteins (yellow).

The analysis of Tfe2-regulated proteins was carried out using Cytoscape applying the String network for *S. cerevisiae* ^29^ (adjusted P-value ≤0.05, fold change >2, Table S3). The early response comprised of 86 proteins significantly altered after 60 min of Tfe2 induction, the majority of which were downregulated. Only three proteins were increased in abundance, namely Gal1, Gal7 and Gal10, which are members of the tightly regulated galactokinase regulon^30^. Differentially regulated proteins were processed by functional enrichment analysis using a Markov Cluster algorithm (MCL). The most prominent early response MCL cluster down-regulated upon Tfe2-activity encompassed proteins involved in amino acid transmembrane transport activity, followed by cell wall proteins and proteins of the sno(s)RNA containing ribonucleoprotein complex (Fig. 2e). The adaptive response comprised of 473 proteins significantly altered after 6h of Tfe2 induction, the majority of which were downregulated. The main MCL cluster included 43 proteins of the ribosome biogenesis and ribonucleoprotein complex, indicating a sustained reduction in the abundance of the translation machinery. Specific to the adaptive response were proteins associated with membrane function, fusion and trafficking, including sterol and sphingolipid biosynthetic processes, intracellular distribution of mitochondria, the mitochondrial envelope and intermembrane space, proteins of the clathrin vesicle coat, the SNARE complex, the ERAD pathway and the Ub-dependent protein transport to vacuole via the MVB sorting pathway (Figure 2f).

These early and adaptive signatures of *S. cerevisiae* responses upon Tfe2 induction directed subsequent investigations to dissect its mode of action and fungal responses to this structurally novel antifungal effector.

### Tfe2 induces amino acid transporter degradation via substrate-independent stress induced endocytosis

As the most prominent early response to Tfe2 toxication is the downregulation of amino acid transporters, the basis for this was explored further. In the presence of extracellular amino acids or certain stress-inducing conditions, amino acid transporters are subject to substrate or stress-induced endocytosis, followed by recycling to the membrane via the trans-Golgi network or trafficking to the vacuole for degradation. Endocytosis is induced by E3 ligase Rsp5-dependent ubiquitination, which is mediated by arrestin-like adapter/facilitator proteins^31, 32^.

To determine if Tfe2 induced downregulation of amino acid transporters is via ubiquitination-mediated endocytosis, we chose to study the broad range amino acid transporter Agp1 for two reasons; Agp1 is rapidly downregulated upon Tfe2 expression in *S. cerevisiae* (Fig. 2e), and the Rsp5-mediated endocytosis of this permease is dependent on only one arrestin-like adapter protein, Bul1^33^. To analyse Agp1 cellular localisation and movement we ectopically expressed Agp1-GFP in WT and *bul1Δ* strains, which carried either the empty *GAL1* promoter or the galactose inducible Tfe2 construct. As Agp1 is subject to substrate-induced endocytosis by a broad range of amino acids, cells were grown in synthetic minimal medium without amino acids and supplemented with allantoin as the sole nitrogen source. Consistent with previous reports, under such growth conditions, Agp1 predominantly localised to the plasma membrane in all strains with small fractions situated on the ER and in the vacuole, with visualisation of the latter carried out by pre-treatment with the membrane stain FM4-64 (Fig. 3). In WT cells, as expected, addition of alanine to the media strongly induced endocytosis of Agp1, resulting in depletion of the fluorescence signal at the plasma membrane and its movement into vesicles and the vacuole after 1 h. After 3 h growth on alanine this was seen to partially re-localise to the ER and plasma membrane, likely due to synthesis of new Agp1 protein. Induction of Tfe2 also resulted in a rapid shuttling of Agp1 from the plasma membrane into vesicular structures and the vacuole after 1 h (Fig. 3). Strikingly however, this was in the absence of alanine, thus illustrating that Tfe2 drives substrate-independent endocytosis of Agp1. Furthermore, Tfe2-induced endocytosis appeared to be more intense compared to the amino acid driven endocytosis in WT cells, as seen by the increased uptake of FM4-64 into the vacuolar membrane both at 1 and 3 h post-treatment. The Tfe2 mediated endocytosis of Agp1 observed in wild-type cells, was inhibited in *bul1Δ* cells (Fig. 3), indicating that arrestin-mediated ubiquitination of the amino acid transporter was key.

**Figure 3.**
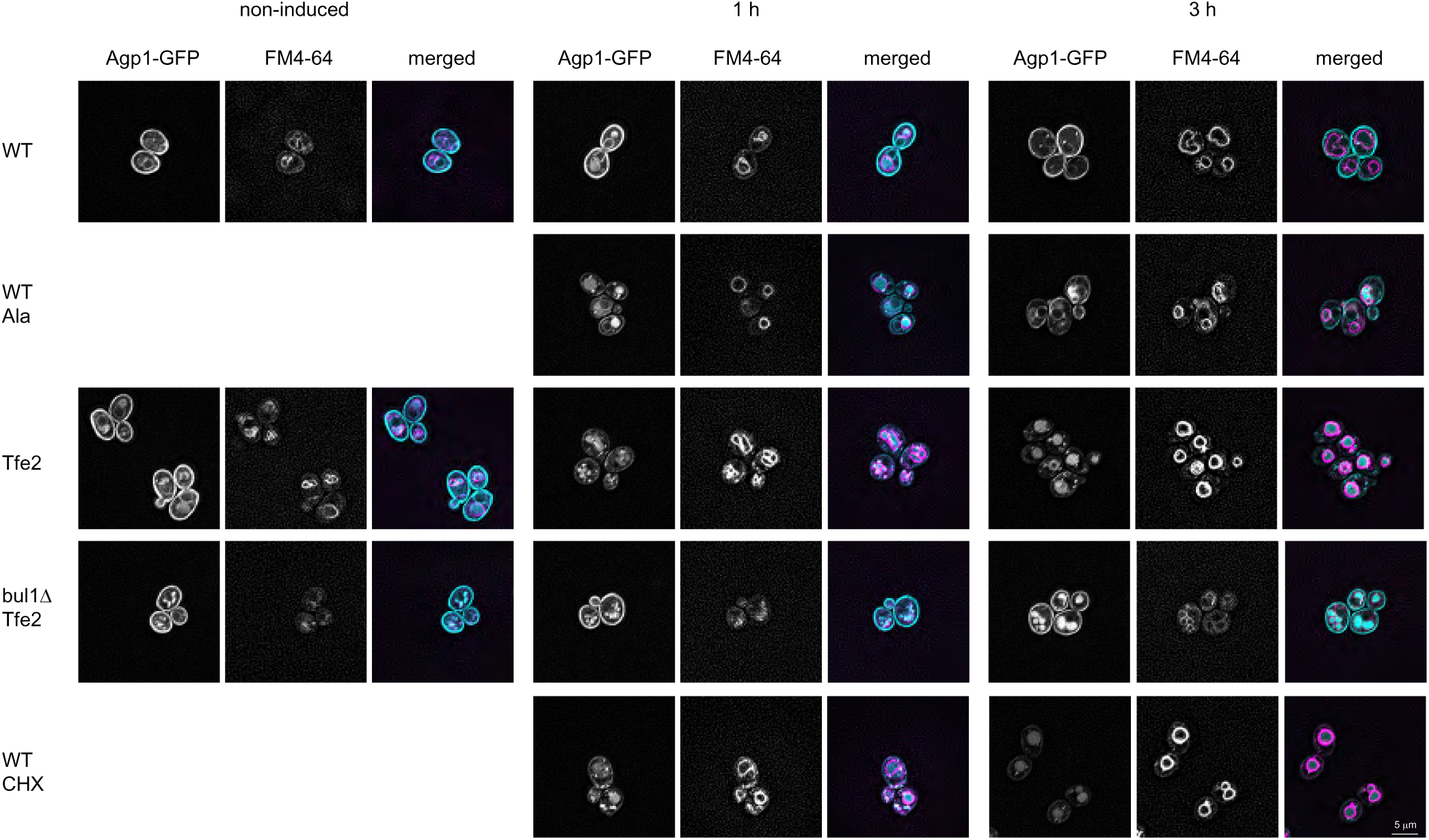
Tfe2-mediated endocytosis of the broad range amino acid transporter Agp1 is substrate-independent requiring the Bul1 arrestin. Localisation of Agp1-GFP by fluorescence microscopy was carried out using the *S. cerevisiae* K699 WT and *bul1Δ* strains carrying either the chromosomally integrated empty GAL-promoter (WT) or the GAL-inducible Tfe2 construct (Tfe2, *bul1*Δ Tfe2). Strains were grown in synthetic minimal medium without amino acids and ammonium sulphate, supplemented with 2% raffinose and 0.1% allantoin as sole nitrogen source. FM4-64 (5 μg/ml) was also added 1 hour prior to the first time point to allow visualisation of the vacuoles followed by culturing of all strains and conditions in the dark. WT and Tfe2 strains were imaged after treatment with 1 % galactose after 0 (non-induced), 1 and 3 h in the respective strains, with WT cells also treated, where indicated, with 10 mM alanine (Ala) or 25 μg/ml cycloheximide (CHX). Cells were imaged using a Zeiss Axiocam Imager.M2 with a Zeiss Plan APOCHROMAT 63x/1.40 Oil Ph3 objective, excitation/emission filters of 450-490/500-550 nm for Agp1-GFP and 538-562/570-640 nm for FM4-64. Widefield image data were deconvolved using Huygens with classic MLE and automatic signal to noise ratio estimation and processed in ImageJ 1.54F keeping parameters constant for all images. GFP and FM4-64 signals in the overlay image were re-coloured in cyan and magenta, respectively, for easy visualisation.

These findings posed the question of how Tfe2 intoxication drives the substrate-independent, ubiquitin-dependent, endocytosis of amino acid transporters? Notably, similar substrate-independent, ubiquitin-dependent, endocytosis has previously been described for the arginine transporter Can1 in response to the translation inhibitor cycloheximide^34^. Thus, we treated Agp1-GFP expressing cells with cycloheximide, and significantly found that this resulted in the same induction of substrate-independent endocytosis of Agp1 as seen upon Tfe2-induction (Fig. 3). Furthermore, in contrast to alanine-treated WT cells, we could not detect any Agp1-GFP signal at the plasma membrane or ER after prolonged Tfe2 induction or cycloheximide treatment, suggesting inhibition of new protein synthesis in response to both stressors.

### Tfe2 leads to an overall increase in cellular amino acid availability and TORC1 hyperactivation

As Tfe2 drives substrate-independent endocytosis of amino acid transporters in a similar manner to cycloheximide, we asked if the mechanism underpinning this was similar. Cycloheximide, due to its ribosome stalling activity resulting in translation inhibition, drives an increase in the intracellular pool of free amino acids, which in turn hyperactivates the TORC1 kinase complex, eliciting the rapid uptake of amino acid transporters^35^. Thus, we set out to investigate if Tfe2 activity similarly triggers an increase in the intracellular pool of free amino acids, and if so whether Tfe2 also drives the hyperactivation of TORC1.

Analysis of the amino acid profiles of WT, Tfe2-intoxicated and cycloheximide-treated cells revealed a remarkable overlap in the profiles following cycloheximide or Tfe2 activity. Most of the 20 essential amino acids were significantly more abundant 30 and 60 min after treatment with cycloheximide, or following induction of Tfe2 for 30 or 60 min (Fig. 4a). A few amino acids, including aspartic acid, threonine, proline and methionine, were downregulated following Tfe2 expression or cycloheximide treatment, but again a striking overlap in profiles was seen. Due to Tfe2 driving the same changes in free amino acids as cycloheximide, we next investigated whether Tfe2 similarly drives the hyperactivation of TORC1, by analysing phosphorylation of one of its key substrates, the kinase Sch9^36^. Analysis of the mobility of Sch9 after Tfe2-induction, or cycloheximide treatment, revealed a second band of lower mobility indicative of post-translational modification. Consistent with this, immunoblots using antibodies to detect TORC1-specific Sch9-Thr^737^ phosphorylation showed low level phosphorylation of Sch9-Thr^737^ in the WT and the Tfe2 strain under non-inducing conditions with the appearance of a second band after 30 min of Tfe2 induction or CHX treatment (Fig. 4b). Significantly, both Tfe2 and cycloheximide-induced lower mobility forms of Sch9 were prevented if cells were simultaneously treated with the TORC1 inhibitor rapamycin (Fig. 4c). Collectively, these experiments show that, like cycloheximide, Tfe2 activity results in an increase in the intracellular levels of many essential amino acids and the hyperactivation of the TORC1 kinase, as evidenced by the rapamycin-sensitive phosphorylation of the Sch9 kinase.

**Figure 4.**
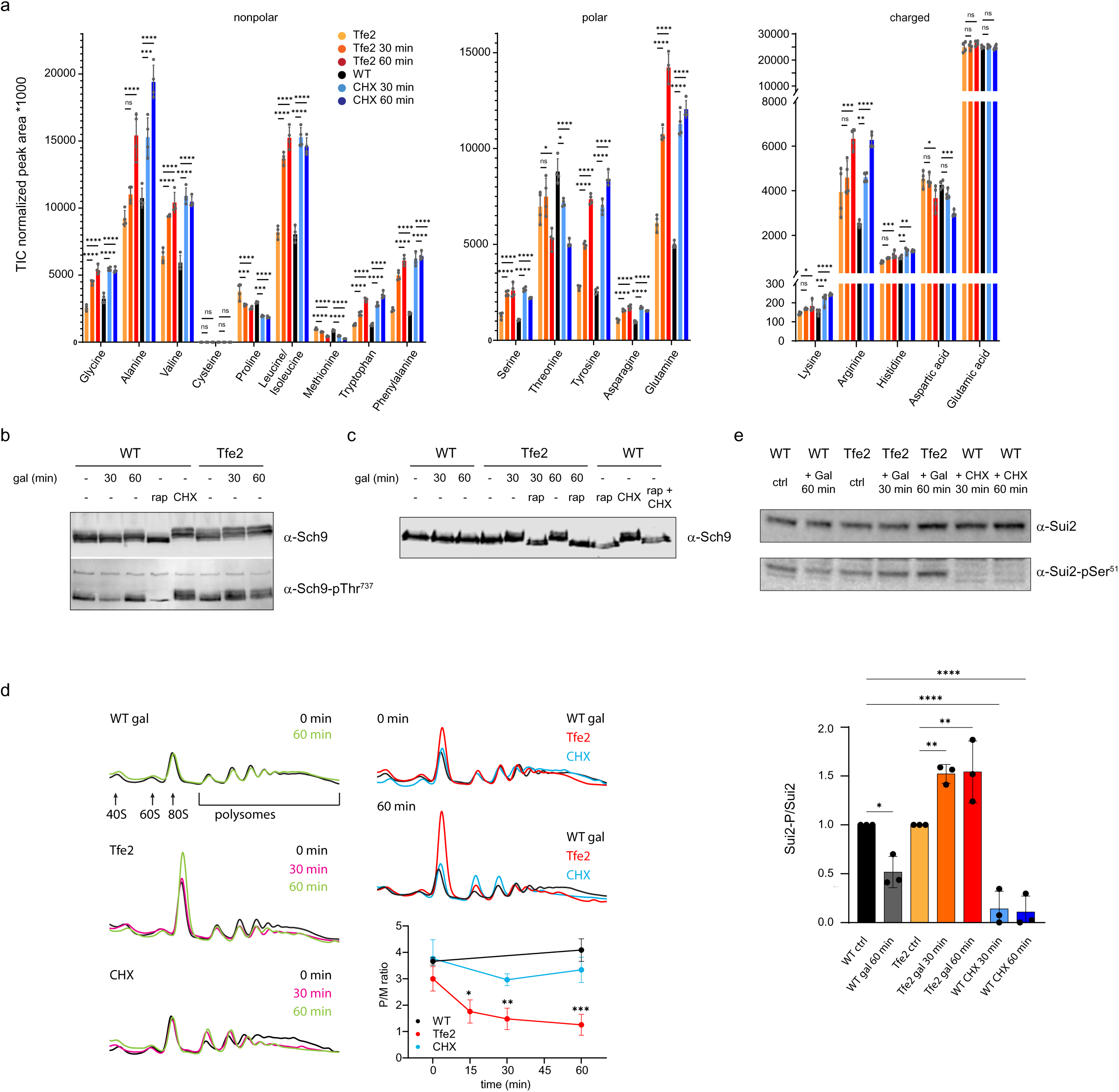
Comparison of Tfe2 intoxication and cycloheximide treatment on cellular amino acid accumulation, TORC1 hyperactivation, and protein translation inhibition. **(a)** Abundance of indicated amino acids in *S. cerevisiae* K699 WT strains carrying either the chromosomally integrated empty GAL-promoter (WT) or the GAL-inducible Tfe2 construct before (WT, black; Tfe2, yellow) and after 30 and 60 min of induction with 1 % galactose (Tfe2, orange) or 25 µg/ml cycloheximide (WT, blue), presented as total ion chromatogram (TIC) normalised peak areas. Data are presented as mean ± SD with individual data points overlaid (n=4 biological replicates; **** P<0.0001, *** P<0.001, ** P<0.01, * P<0.05, ns not significant; one-way ANOVA of the respective amino acid over all strains and conditions with Tukey’s test). **(b,c)** Immunoblots of the TORC1 substrate Sch9, in control (WT) and Tfe2-harbouring strains after induction with 1 % galactose at indicated timepoints and/or treatment with 25 µg/ml cycloheximide and/or 200 ng/ml rapamycin for 20 min. Lysates were analysed following western blotting using anti-Sch9 (α-Sch9) and anti-Sch9-pThr^737^ (α-Sch9-P) antibodies Rapamycin treatment abolished the slower migrating forms of Sch9 induced by both Tfe2 intoxication or cycloheximide treatment. **(d)** Polyribosome profiles of control (WT) and Tfe2-harbouring strains. Cells were harvested before and after induction with 1 % galactose or treatment with 25 µg/ml cycloheximide for 30 or 60 min. Shown are overlays of representative profiles of each strain/condition at different timepoints (left panels) and all strains/conditions at timepoints 0 and 60 min (upper and middle right panels). 40S, 60S, 80S and polysome fractions are indicated on the wildtype profile overlay. Relative levels of polysomes/monosomes per strain and condition are presented as mean ± SD (lower right panel, n=3 biological replicates; *** P<0.001, ** P<0.01, * P<0.05; one-way ANOVA with Šídák’s test). **(e)** Immunoblot of the eIF2α subunit Sui2 in control (WT) and Tfe2-harbouring strains after induction with 1 % galactose or treatment with 25 µg/ml cycloheximide at indicated timepoints. Lysates were analysed using anti-Sui2 (α-Sui2) and anti-elF2α-pSer^51^ (α-Sui2-P) antibodies (top panel). Relative levels of Sui2/Sui2-P are presented as mean ± SD with individual data points overlaid (n=3 biological replicates; **** P<0.0001, ** P<0.01, * P<0.05; one-way ANOVA with Šídák’s test; bottom panel)

### Tfe2 reduces global protein synthesis via inhibition of translation initiation

Following on from the findings above, we asked if Tfe2, like cycloheximide, inhibits protein translation. To facilitate this we performed polyribosome profiling on *S. cerevisiae* WT cells, and cells following cycloheximide treatment or Tfe2-induction. Polysomes are ribosomes that are actively translating mRNAs. They can be separated on sucrose density gradients and quantified by absorbance measurement at 253 nm. As can be seen for WT cells this allows for the identification of the 40S and 60S ribosomal subunits, as well as monosomes (80S ribosomes) and polysomes^37^ (Fig. 4d). Treatment with cycloheximide gave rise to a distinct polyribosome profile with an unaffected 80S monosome peak and an increase in the di- and trisome peaks, a profile indicative of ribosome stalling, consistent with impairment of translocation^38^. Interestingly, however, in the Tfe2-harbouring strain a strong increase in the 80S monosome peak was already apparent at time-point 0, before the addition of galactose, which is likely due to the leakiness of the *GAL1* promoter when cells are grown on raffinose. After 60 min of Tfe2-expression the 80S peak increased further, with a slight decrease in the polysome peak area. The accumulation of ribosomes in the 80S peak of a sucrose gradient is indicative of decreased translation initiation, and is reflected in the decrease of the polysome over monosome ratio over time when compared to WT cells^39^ (Fig. 4d). The prominent increase in monosomes seen in Tfe2-expressing cells, is further in agreement with analysis of the ribosome density by transmission electron microscopy, with a 1.7-fold increase of ribosomes per area of cytosol upon Tfe2-intoxication (fraction of ribosomes per area of cytosol WT 0.2053 %, Tfe2 0.3524 %).

To interrogate the Tfe2-specific inhibition of translation initiation further, we analysed the phosphorylation profile of Sui2, the α-subunit of eukaryotic translation initiation factor 2 (elF2α). Phosphorylation of eIF2α is induced in response to a diverse range of environmental challenges, including amino acid starvation, and ER, osmotic or oxidative stress^37, 40^, leading to the inhibition of translation. In WT cells, using a Sui2-P specific antibody, we noted a two-fold decrease in Sui2-P following switching from raffinose to the preferred carbon source galactose (Fig. 4e), likely due to an increase in growth rate. In contrast, upon Tfe2 induction we detected a modest increase of Sui2-P signal (1.5-fold) which was directly opposite to that seen with cycloheximide treatment which severely compromised Sui2-P (Fig. 4e). Taken together, the observations that Tfe2 activity and cycloheximide result in different polysome profiles and have opposite effects on elF2α-phosphorylation, support a model where Tfe2 and cycloheximide lead to translation inhibition using distinct modes of action.

### Tfe2 drives changes to the cellular lipid profile of yeast cells

After characterising the initial cellular responses to Tfe2, we next analysed longer term responses to this effector informed by the proteomics data obtained following 6 h induction. We were particularly interested in the observation that Tfe2 intoxication led to the reduction of multiple enzymes involved in lipid metabolism, especially the sphingolipid (SL) biosynthetic pathway (Fig. 2f), as previous work has linked TORC1 activity to the regulation of sphingolipid synthesis in *S. cerevisiae*^41^. Sphingolipids play key roles in fungal physiology and are important virulence determinants in pathogenic fungi, contributing to plasma membrane stability, acting as membrane anchors for proteins and contributing to microdomains which drive endocytosis, nutrient transport and pH maintenance^42, 43^. Whilst enzymes responsible for synthesis of the complex sphingolipids inositol phosphorylceramide (IPC; Aur1), (MIPC; Sur2) and mannosyl-di-IPC (M(IP)_2_C; Ipt1), were found to be less abundant after 1 h of Tfe2 induction, prolonged exposure of 6h resulted in a further decrease of these proteins as well as downregulation of additional proteins in the sphingolipid synthesis pathway (Fig. 5a), suggesting interference of the whole pathway.

**Figure 5.**
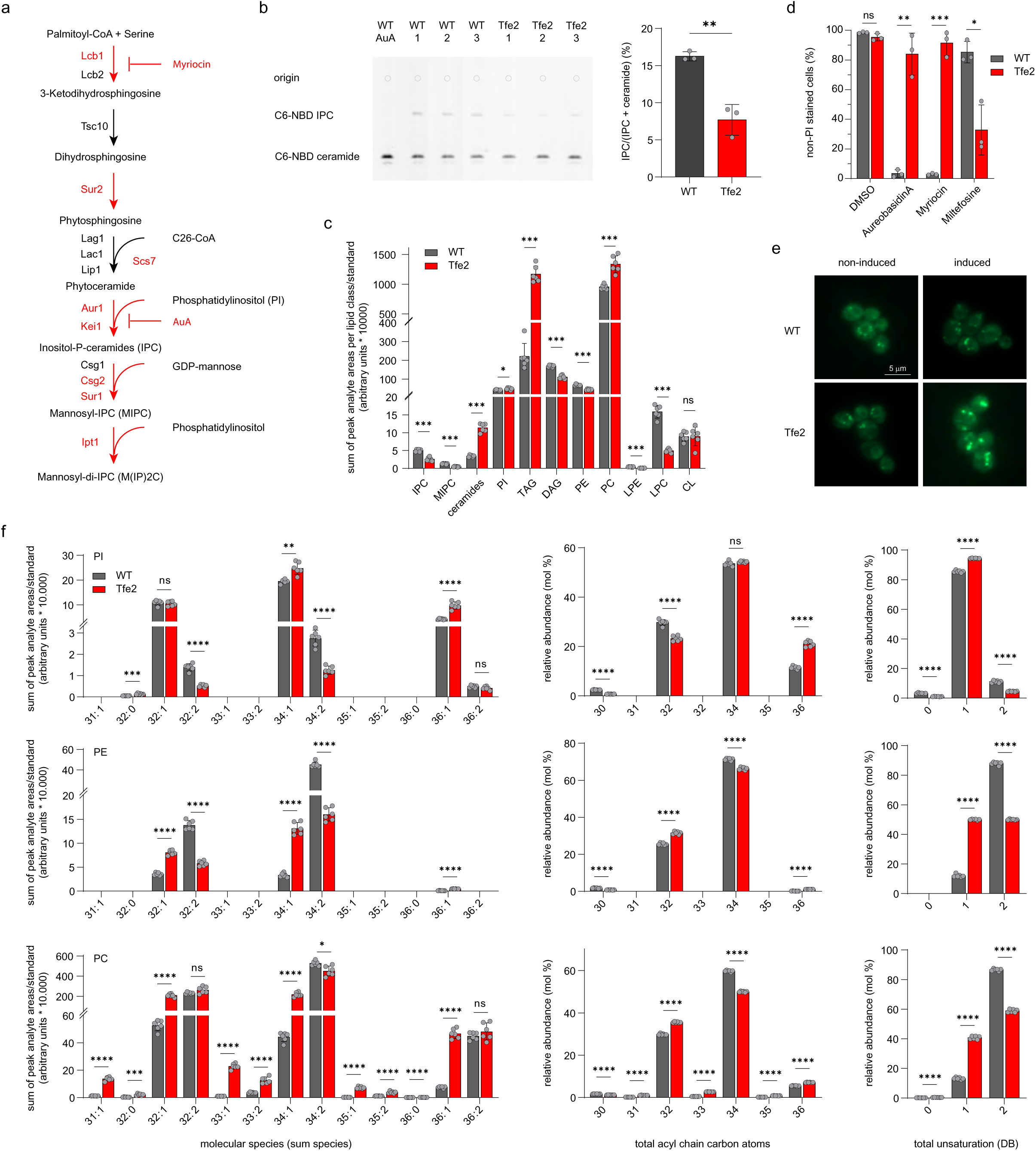
Tfe2 drives changes in the lipid profile of yeast cells. **(a)** Schematic of the sphingolipid synthesis pathway in yeast with Tfe2-downregulated enzymes in red. Point of action of the sphingolipid synthesis inhibitors Myriocin and AureobasidinA (AuA) are indicated. **(b)** Thin layer chromatogram of lipid extracts from control (WT) and Tfe2-induced yeast cells grown in the presence of C6-NBD ceramide. Origin of samples on top as indicated. Incubation of control (WT) cells with 0.25 µg/ml AuA was used as control for Aur1 inhibition. Relative levels of C6-NBD-IPC over combined levels of C6-NBD-IPC and C6-IPC-ceramide are shown to the right, presented as mean ± SD with individual data points overlaid (n=3 biological replicates; ** P<0.01; unpaired t-test). **(c)** Changes in lipid class composition to Tfe2-intoxication depicted by sums of lipid class molecular species of control (WT) and Tfe2-induced cells as determined by lipid chromatography mass spectrometry analysis. Data are presented as mean ± SD with individual data points overlaid (n=6 biological replicates; *** P<0.001, ** P<0.01, * P<0.05; ns not significant; multiple unpaired t-tests with Holm-Šídák’s test). IPC – inositolphosporyl ceramide, MIPC – mannosyl-IPC, PI – phosphatidylinositol, TAG – triacylglyceride, DAG – diacylglyceride, PE - phosphatidylethanolamine, PC – phosphatidylcholine, LPE – lyso-PE, LPC – lyso-PC, CL – cardiolipin. **(d)** Impact of Tfe2 toxication on susceptibility to lipid-targeting antifungal drugs. Control and Tfe2-cells were grown in the presence of 1 % galactose for 5h30 and treated with 500 ng/ml AureubasidinA, 40 µg/ml Myriocin and 2.5 µg/ml Miltefosine with DMSO as control for 16 hours. Cell survival was calculated by staining with propidium iodide (PI) and FACS analysis. Data are presented as mean ± SD with individual data points overlaid (n=3 biological replicates; *** P<0.001, ** P<0.01, * P<0.05; ns not significant; multiple unpaired t-tests with Holm-Šídák’s test). **(e)** Fluorescence microscopy of lipid droplets in control (WT) and Tfe2-harbouring yeast cells before (non-induced) and after 90 min induction with 1 % galactose (induced) using 5 µg/ml BODIPY 493/503. Scale bar 5 µm. **(f)** Impact of Tfe2 toxication on the composition of phospholipids, shown as sum-species composition, total carbon atom length and number of double bonds of PI (upper panel), PE (middle panel) and PC (lower panel). Total carbon atom length and number of double bonds are presented as the sum of the number of carbon atoms and the number of double bonds in both acyl chains, respectively. Data shown are for molecular species with total carbon lengths of C31 to C36 only. Data are presented as mean ± SD with individual data points overlaid (n=6 biological replicates; *** P<0.001, ** P<0.01, * P<0.05, ns not significant; multiple unpaired t-tests with Holm-Šídák’s test).

To validate if the proteomic downregulation of sphingolipid biosynthesis enzymes by Tfe2 correlated with a measurable decrease in enzymatic activity, we determined the activity of the IPC synthase Aur1. Quantification of IPC production was carried out by thin layer chromatography using membrane permeable fluorescent-labelled ceramide (C6-NBD ceramide) as precursor^44^ and the Aur1 inhibitor Aureobasidin A (AuA) as control treatment (Fig. 5b). After 6 h of Tfe2 induction, IPC levels were reduced 2.1-fold compared to WT cells indicating that the Tfe2 mediated decrease in Aur1 protein levels does corelate with a decrease in Aur1 enzymatic activity.

Having confirmed the direct effect of Tfe2 on IPC levels, we asked if Tfe2 specifically impacts on sphingolipid synthesis or induces more wide-ranging changes to the lipid profile of the yeast cell, as also suggested by the proteomics analysis (Fig. 2f). WT and Tfe2-intoxicated cells were subjected to liquid chromatography-coupled mass spectrometry-based analysis which allowed for the identification of 11 classes of lipids (Fig. 5c, Table S4). Importantly, both IPC and MIPC sphingolipids were reduced in Tfe2-intoxicated cells by 2- and 2.8-fold, respectively, consistent with the proteomics data showing downregulation of sphingolipid biosynthesis enzymes. Interestingly, precursors of IPC biosynthesis, ceramides and phosphatidylinositol (PI), were increased by 3.3- and 1.2-fold, respectively, suggesting inhibition of IPC to be the driving factor in the observed decrease in complex sphingolipids. This is in agreement with the proteomics dataset which did not detect changes in the abundance of the ceramide synthase subunits Lag1, Lac1 and Lip1.

The Tfe2-mediated decrease in sphingolipid levels prompted an exploration as to whether Tfe2 intoxication impacted on resistance to lipid-targeting antifungals. Control and Tfe2-expressing cells were grown in the presence of galactose for 5h30 prior to treatment with the sphingolipid targeting drugs Aureobasidin A and myriocin, which inhibits serine palmitoyltransferase that catalyses the first step in sphingosine biosynthesis (Fig. 5a), and the phospholipid mimicking drug miltefosine. Cell survival was analysed by FACs analysis of propidium iodide (PI) stained cells (Fig. 5d) and colony forming unit (cfu) counts (Fig. S2). This clearly indicated that Tfe2 intoxication results in resistance to both sphingolipid targeting drugs compared to WT cells, but is more susceptible to the alkylphosphocholine miltefosine.

From the lipid profiling, it was clear that Tfe2 impacted on other lipid classes in addition to sphingolipids. The largest overall change was seen for the neutral lipid class of triacylglycerols (TAG), whose levels increased 5.3-fold upon expression of Tfe2. TAGs have been reported to accumulate at later growth stages, serving as easy access reservoirs of fatty acids, and are stored in lipid droplets^45^. Microscopic fluorescence analysis using the neutral lipid stain Bodipy revealed an increase in the number and size of lipid droplets upon Tfe2 expression (Fig. 5e), consistent with the large increase in TAGs.

Tfe2-intoxication also impacted on the most abundant class of lipids, the glycerophospholipids, resulting in a reduction in phosphatidylethanolamine (PE) levels (1.5-fold), and elevated levels of phosphatidylcholine (PC; 1.4-fold) and PI (1.2-fold; Fig. 5f). For all three phospholipid classes, the majority of lipids consisted of acyl chains totalling 32, 34 and 36 carbon atoms and harbouring a total of 1 or 2 double bonds, which is in accordance to previous findings^46^. The Tfe2-induced change in the lipid species profile of PI resulted in longer acyl chains and a slight increase in mono-unsaturated species as compared to double unsaturated species. This was distinct to the profile changes observed for both PE and PC lipid classes, for which the total carbon chain length decreased and the initial ratio of mono-to di-unsaturated lipids of roughly 10:90 increased to nearly equal distribution upon Tfe2 intoxication. We further detected an increase in odd-numbered carbon chain length species for PC, the most abundant phospholipid in both WT and Tfe2-induced cells. Overall, Tfe2 intoxication resulted in decreased saturation of the acyl chains across all three phospholipid classes, which was also reflected upon analysis of total acyl chain composition (Fig. S3), and longer carbon chains for both PI and PC.

### Tfe2 induces wide-ranging changes to the fungal cell wall and organelle structures

In addition to changes in cellular lipid composition following Tfe2 intoxication, the proteomics analysis revealed that Tfe2 activity potentially drives changes in the cell wall and mitochondria (Fig. 2f). To explore this further, and complete our comprehensive analysis of Tfe2, an overview of cellular structural changes induced upon Tfe2 intoxication was obtained by performing transmission electron microscopy (TEM) of yeast cells. TEM images of chemically fixed cells revealed a general thickening of the cell wall, the appearance of irregular broadened patches specific to the inner cell wall, and major distortion of the nucleus and its ER envelope (Fig. 6a). To further analyse these Tfe2-induced cell wall alterations, the width of the cell wall was quantified using TEM of freeze-substituted cells (Fig. 6b), which revealed an increase in both the inner and outer cell wall layers upon Tfe2-intoxication (Fig. 6c). The inner layer, composed of β-glucan and chitin, determines mechanical flexibility and strength of the cell wall, whilst the outer layer, consisting of glycosylated mannoproteins, determines the biophysical properties of the fungal surface and its interaction with the environment^47^. Subsequently, the relative carbohydrate composition of the cell wall was determined by HIPC analysis. This revealed an increase in both the glucose (β-glucan) and glucosamine (chitin) fractions in expense of the mannose fraction upon expression of Tfe2 (Fig. 6d). Calcofluor white staining of yeast cells confirmed an increase in chitin levels upon Tfe2 intoxication, localized as patches to the cell wall, likely representing the broadened inner cell wall patches seen by TEM imaging (Fig. S4).

**Figure 6.**
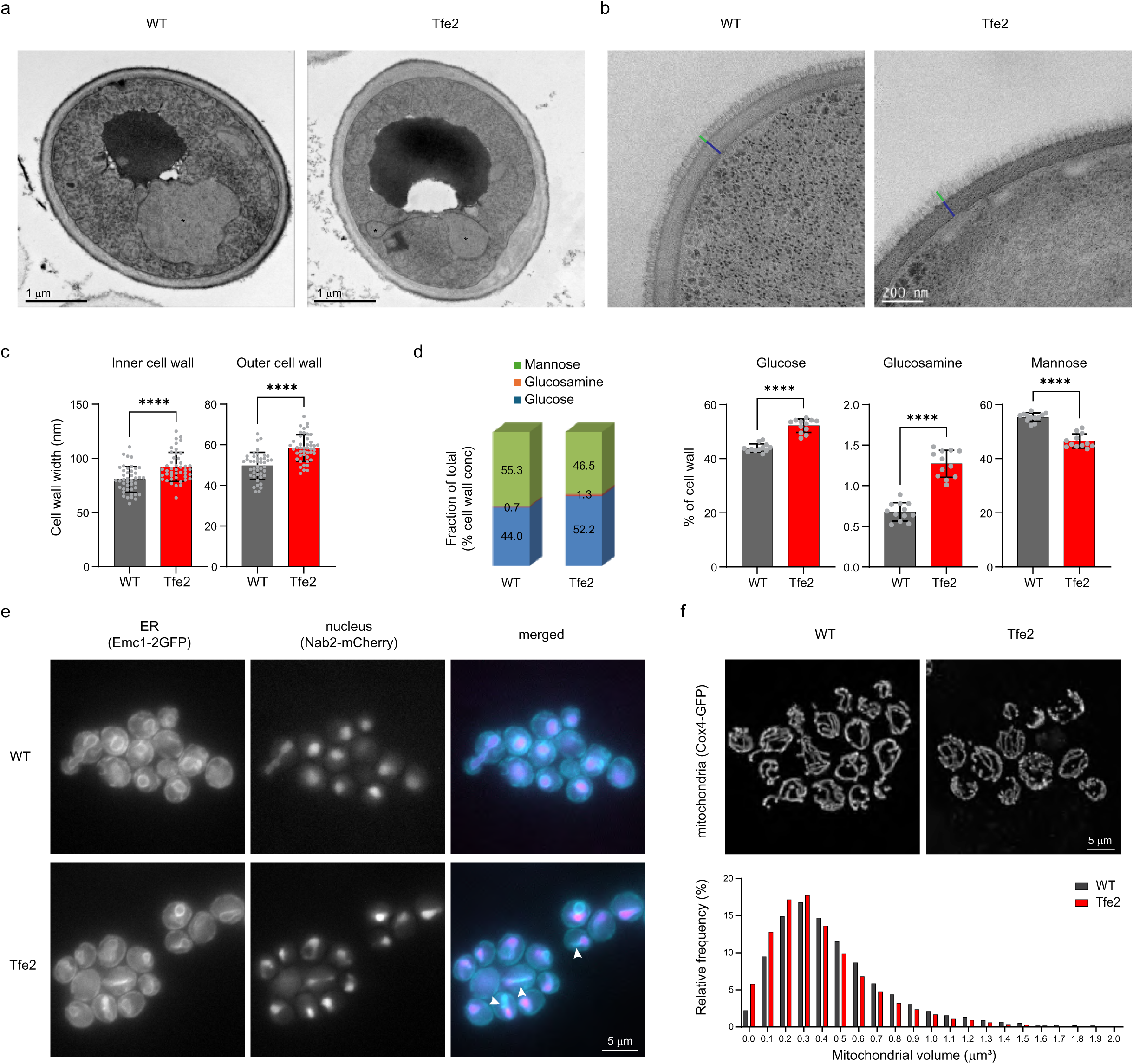
Tfe2 intoxication changes the fungal cell wall and organelle structures. **(a)** Electron micrographs of control (WT) and Tfe2-expressing whole yeast cells harvested by glutaraldehyde/paraformaldehyde fixation followed by potassium permanganate staining. Nuclear (sub-)structures are indicated by asterisks. Scale bare 1 µm. **(b)** Electron micrographs of control (WT) and Tfe2-expressing yeast cell wall harvested by cryofixation and freeze substitution with osmium tetroxide. Inner (blue) and outer (green) cell wall sections are indicated per cell. Scale bar 200 nm. **(c)** Changes in both the inner and outer cell wall width of control (WT) and Tfe2-expressing cells harvested by cryofixation and freeze substitution with osmium tetroxide. Data are presented as mean ± SD with individual data points overlaid (n = 43 (WT) and 45 (Tfe2); **** P<0.0001; unpaired t-test). **(d)** Carbohydrate cell wall composition of control (WT) and Tfe2-inducing strains. The amount of glucosamine (from chitin), glucose (from glucan) and mannose (from mannan) in the cell wall was determined by HPIC. Data are presented as fraction of the total cell wall showing the mean ± SD with individual data points overlaid (n=12, 3 biological replicates measured in quadruplicate; **** P<0.0001; unpaired t-test). **(e)** Fluorescence microscopy of ER (cyan) and nuclear (magenta) structures in yeast control (WT) and Tfe2-inducing strains carrying the chromosomally integrated organelle-specific fluorophore-tagged proteins Emc1-2GFP (ER) and Nab2-mCherry (nucleus). **(f)** Fluorescence microscopy of the mitochondrial network in WT and Tfe2-inducing strains carrying the chromosomally integrated mitochondrial fluorophore-tagged protein Cox4-GFP. Presented are the relative frequency of volumes of distinct mitochondrial units between 0 and 2 µm^3^ per strain (n=15 fields of view with a minimum of 15000 units per strain).

The Tfe2-induced changes in organelle structures were explored further. The defects in nuclear and ER structures, observed in the TEM images (Fig. 6a), were investigated by live-cell microscopy of nucleus and ER-specific fluorophore-tagged marker proteins (Nab2-mCherry, Emc1-2GFP^48^). This confirmed loss of the spherical structure of the nuclear envelope and the appearance of bi-lobed nuclear structures as early as 30 min after induction (Fig. 6e, Fig. S5). Significant changes in ER structures were also clear upon Tfe2 induction. The nuclear ER ring which is continuous with the nuclear envelope is significantly deformed following Tfe2 induction, and in some cells shows separation from nuclear structures (Fig. 6e). Following on from the proteomics data showing Tfe2-induced down regulation of proteins linked to the intracellular distribution of mitochondria, mitochondrial inner membrane space, and the mitochondrial envelope, we examined mitochondrial structures following Tfe2 intoxication of yeast cells. Tfe2 intoxication resulted in significant fragmentation of the mitochondrial network visualised using the mitochondrial reporter protein Cox4-GFP. This was confirmed by volumetric measurement of 3D images, which showed an increase in smaller volume entities upon Tfe2-intoxication as compared to the WT (Fig. 6f). Structural changes to both the nuclear ER envelope and the mitochondrial network could also be detected in yeast cells following co-culturing with *S. marcescens*, which was both T6SS- and Tfe2-dependent (Fig. S6), confirming that such alterations were conserved upon T6SS-mediated delivery of Tfe2.

Taken together the proteomics data, coupled with electron and fluorescent microscopy approaches, revealed that Tfe2 intoxication of yeast cells results in significant changes to the cell wall and the structure of intracellular organelles.

## DISCUSSION

In this study we reveal that Tfe2 is an antifungal effector that adopts a novel structure and triggers the rapid inhibition of protein translation initiation (Fig. 7). This in turn drives an increase in the free pool of intracellular amino acids, hyperactivation of the TORC1 kinase, and rapid endocytosis of amino acid and other transporters (Fig. 7). Adaptative fungal responses to this effector include significant lipid remodelling and dramatic cell wall and organelle structural alterations (Fig.7). Collectively these findings underpin the potent activity of Tfe2 to inhibit the growth of both pathogenic and model yeasts^9^ and reveal a previously undescribed mode-of-action of a T6SS-elicited effector.

**Figure 7.**
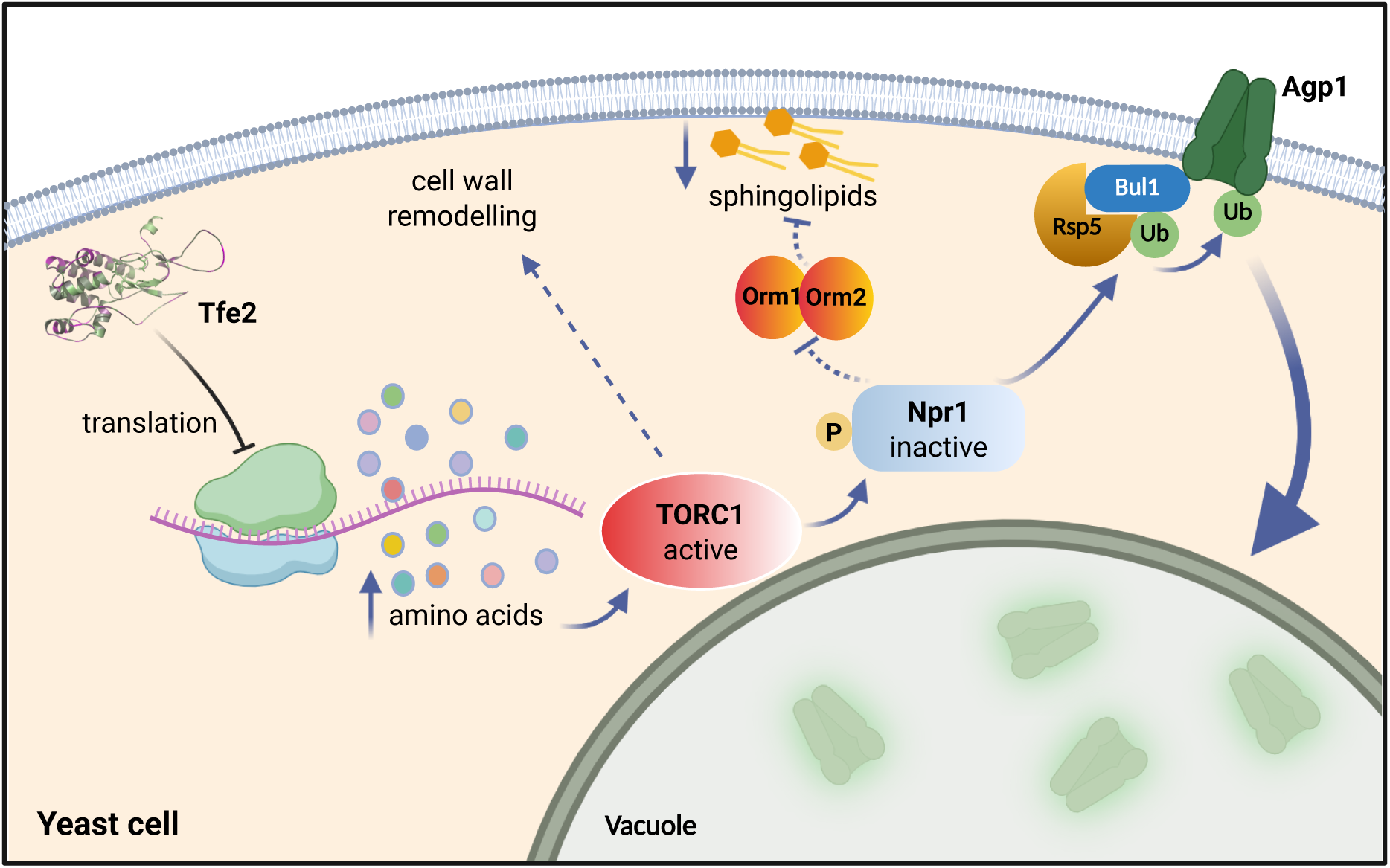
Model underpinning the antifungal activity of the *S. marcescens* T6SS antifungal effector Tfe2. The *S. marcescens* T6SS injects the antifungal effector Tfe2 into neighbouring yeast cells where it quickly inhibits protein translation, resulting in an increase in intracellular amino acids that hyperactivates the TORC1 kinase. Hyperactivation of TORC1 results in the phosphorylation and inhibition of the Npr1 kinase substrate, which can no longer phosphorylate arrestin adaptor proteins such as Bul1. In the unphosphorylated form, Bul1 can recruit the Rsp5 ubiquitin ligase to amino acid transporters such as Agp1, resulting in ubiquitination which drives endocytosis to the vacuole and degradation. Based on previous studies, hyperactivation of TORC1 may also underlie the Tfe2-mediated reduction in sphingolipids, through preventing the Npr1-mediated activation of the sphingolipid regulators Orm1/2. In addition, hyperactive TORC1 has been shown to result in cell wall modelling which is also seen upon Tfe2 intoxication of *S. cerevisiae*. Created with BioRender.com.

Our structural and bioinformatic analysis indicate that Tfe2 is a conserved effector protein adopting a novel fold. It therefore represents a previously uncharacterised effector family, which is found across the bacterial classes of Alpha-, Beta- and Gammaproteobacteria^22^. Whilst our structural and mutagenesis data support an enzymatic activity via an acid-base hydrolysis mechanism, we could not confirm its cellular target due to lack of purified protein for enzymatic analysis, or its precise cellular localisation or interaction partners due to extremely low expression levels upon expression in *S. cerevisiae* (Fig. S1). The lack of predictable functional motifs, and inability to purify Tfe2 for enzymatic assays, prompted a non-biased proteomic analysis to accurately capture the immediate and longer-term responses of *S. cerevisiae* cells to Tfe2 intoxication. This dataset resulted in 2007 ANOVA positive proteins, of which 86 proteins formed the early cellular response to Tfe2, and 473 proteins constituted the late response, the majority of which were downregulated. This approach proved extremely informative as the Tfe2-driven proteomic changes directed a series of experiments that significantly increased our knowledge of mode-of-action and fungal responses to this effector (Fig. 7).

Previous in-competition proteomics following extended co-culture of *S. marcescens* with *C. albicans* cells revealed that Tfe2 impacts on amino acid transporter turnover^9^. In this study, we show that downregulation of plasma membrane amino acid transporters is the most prominent early response of *S. cerevisiae* cells to Tfe2 activity. Regulation of amino acid transporters in *S. cerevisiae* is complex and occurs both at the level of transcription and through the trafficking of transporters to and from specific membranes^49^. Here we find that Tfe2 activity drives the rapid endocytosis of amino acid transporters from the plasma membrane to the vacuole. However, it is noteworthy that two transcriptional regulators of amino acid transporter genes, Gcn4 regulated by the general amino acid control (GAAC) pathway, and Stp1 regulated by Ssy1-Ptr3-Ssy5 (SPS) signalling^49^, were also downregulated in the early proteome of Tfe2-induced cells. Thus, for some transporters there may also be a level of transcriptional regulation. However, we think it unlikely that transcription regulation is a major determinant underpinning Tfe2-mediated reduction in nutrient transporter levels, as a number of membrane transporters not under the control of Gcn4 or Stp1 regulation, including the histidine permease Hip1, the manganese transporter Smf1, and the galactose permease Gal2, were also reduced upon Tfe2 intoxication.

Endocytosis of amino acid transporters is generally observed in the presence of amino acid substrates, and is triggered by the ubiquitination of specific Lys residues which in *S. cerevisiae* is mostly mediated by the Rsp5 ubiquitin ligase^50^. Recruitment of Rsp5 to transporters is mediated by alpha arrestin-like adaptor proteins^51^, which are activated following amino acid transport into the cell. Specifically, amino acid import stimulates the TORC1 kinase which leads to the phosphorylation and inactivation of its kinase substrate, nitrogen permease reactivator 1 (Npr1), a negative regulator of arrestins^52^. Notably, all Tfe2-targeted plasma membrane transporters identified in this study, including the non-amino acid transporters, Smf1^53^ and Gal2^54^, are known to be ubiquitinated by the Rsp5 ubiquitin ligase^55^. Furthermore, by following the cellular localisation of the broad amino acid transporter Agp1, we demonstrate that Tfe2 drives the rapid substrate-independent endocytosis of Agp1 in a mechanism dependent on the Bul1 arrestin. Similar substrate-independent endocytosis/degradation of amino acid transporters in *S. cerevisiae* has been reported for a number of stress agents^56^, including the translation inhibitor cycloheximide^34, 52^. Cycloheximide, by blocking protein translation, is predicted to increase intracellular amino acid levels, resulting in hyperactivation of TORC1, and inhibition of the Npr1 kinase, allowing for activation of arrestin activity^34^. Significantly, we demonstrate in this study that Tfe2 is also a potent inhibitor of translation, resulting in the same increase in free amino acid levels, and hyperactivation of TORC1, as seen with cycloheximide. This likely underpins the identical arrestin-mediated endocytosis of Agp1 observed in this study upon cycloheximide treatment or Tfe2-induction.

The observation that Tfe2 inhibits protein translation is a major breakthrough in this study. Cycloheximide blocks translation elongation by binding to the E site of the large ribosomal subunit, resulting in ribosome stalling^38, 57^. However, here we show Tfe2 inhibits translation initiation as polysome profiling revealed Tfe2-driven increases in monosomes. Stress-induced phosphorylation of elongation initiation factor 2 (elF2α) is a common readout of translation initiation inhibition. *S. cerevisiae* has a single elF2α kinase, Gcn2, which is activated by binding uncharged tRNAs during amino acid starvation conditions, as well as starvation-independent activation caused by certain stresses including oxidative stress^37^, or ribosome stalling via the ribosomal P stalk proteins P1/P2^58^. Treatment with cycloheximide resulted in the complete loss of elF2α-phosphorylation, whilst Tfe2-intoxication caused a moderate increase in elF2α-P. As there is an increase in the intracellular amino acid pool in Tfe2 intoxicated cells it is unlikely that uncharged tRNAs are triggering activation of Gcn2. Phosphorylation of eIF2α causes a global inhibition of protein synthesis as well as gene-specific translational activation of *GCN4*^59^. However, as mentioned above, Gcn4 protein levels are strongly downregulated after Tfe2-induction (−1.8-fold at 1h and −2.5-fold at 6h), an observation described previously for replicatively aged cells^60^. Thus, collectively, the data presented here indicate that Tfe2 activity drives the inhibition of protein translation in a mechanism that may involve phosphorylation of eIF2α. Further investigation is needed to link the predicated enzymatic activity of Tfe2 with protein translation inhibition in yeast cells.

Regarding the adaptive responses of yeast cells to prolonged Tfe2 intoxication, a striking finding was the downregulation of many enzymes in the sphingolipid biosynthesis pathway. Enzymatic assays and lipid profiling confirmed that Tfe2 activity drives the downregulation of the sphingolipid biosynthesis enzyme Aur1, and total sphingolipid levels, culminating in resistance to sphingolipid targeting antifungals. Can this response to Tfe2 activity be linked to Tfe2-mediated protein translation inhibition and TORC1 hyperactivation? Notably, previous studies have shown that TORC1 inhibition through rapamycin treatment alleviates the TORC1-inhibition of the Npr1 kinase, which then directly phosphorylates the sphingolipid homeostasis regulators Orm1 and Orm2, promoting the de novo synthesis of complex sphingolipids^41^. Furthermore, cycloheximide treatment, which hyperactivates TORC1, inhibits phosphorylation of Orm1 and 2^41^. This supports a model whereby cycloheximide and, by extension, Tfe2, inhibit sphingolipid synthesis through translation inhibition-induced TORC1 hyperactivation (Fig. 7).

Finally, informed by our proteomic and lipidomics analyses we found that Tfe2 intoxication also caused notable changes in the fungal cell wall, and significant perturbation in the morphology of the ER/nucleus and the induction of mitochondrial fragmentation. Interestingly, the T6SS-elicited effector VgrG4 also drives mitochondrial fragmentation in yeast and mammalian cells, by stimulating calcium influx into the mitochondria^20^. It is possible that Tfe2-induced mitochondrial fragmentation reported here, together with other changes in organelle structures, are due to the significant remodelling of the fungal lipidome. This is likely to effect the lipid composition of membranes which is closely linked to properties including membrane fluidity and curvature, which impact on membrane protein assembly and organellar function^46^. Notably, the largest impact of Tfe2 on the *S. cerevisiae* lipidome was an increase in TAGs which likely underpins the Tfe2-induced surge in lipid droplet formation. This organelle is important for lipid storage, trafficking and homeostasis, and is highly versatile in its ability to interact with nearly all other organelles and can accommodate rapid changes in size and lipid composition^61^. The increase in lipid droplets following Tfe2 intoxication suggests an increased requirement for lipid exchange and storage. Regarding Tfe2-mediated changes in fungal cell wall architecture it is possible that these too may, at least in part, be an adaptation of TORC1 hyperactivation. Previous work revealed that *S. cerevisiae* cells expressing a hyperactive *TOR1^L2134M^* allele resulted in alterations in cell wall architecture^62^, and similar to that seen upon Tfe2 intoxication, exhibited a significant increase in chitin levels. Whatever the underlying mechanism, Tfe2-mediated changes both in the composition and the thickness of the cell wall will dramatically impact the physiological properties of the fungal cell.

Whilst understanding how the T6SS operates as a biological antifungal weapon is of fundamental scientific interest, the identification of dedicated antifungal effectors and determination of their mode-of-action also offers opportunities for developing new antifungal strategies. This field is in its infancy, with only a handful of antifungal effectors being identified to date. Here our observation that the *S. marcescens* antifungal effector Tfe2 drives protein translation inhibition and TORC1 hyperactivation adds a new facet to T6SS-mediated activity against eukaryotic competitors.

## MATERIALS AND METHODS

### Fungal strains, plasmids and culture conditions

Fungal strains and plasmids used in this study are detailed in Table S5. *S. cerevisiae* strains were derived from either K699^63^ or BY4741^64^. Fungal strains were cultured at 30°C in YPDA (10 g/L yeast extract, 20 g/L peptone, 40 mg/L adenine hemisulfate) or synthetic complete media (SC; 6.9 g/L yeast nitrogen base complete with amino acids, with adenine at 100 mg/L) containing 2% glucose, with 20 g/L agar for solid media. For selection of auxotrophic markers, SC media lacking the respective amino acid or nucleotide was used. For induction of the P*_GAL1_* promoter, *S. cerevisiae* cultures were pre-incubated in SC media containing 2% raffinose, followed by induction of gene expression by the inclusion of 1% galactose, unless stated otherwise.

### Tfe2 multiple sequence alignment and AlphaFold modelling

A search for Tfe2 homologues was performed using NCBI BLASTP against the nr database, with inclusion criteria of 25-100% sequence identity and ≥60% query coverage. From the resulting dataset, one representative sequence per species was retained to generate a non-redundant alignment set. Poorly aligned or gapped regions were manually trimmed to focus on the conserved core. Sequences were aligned using the MAFFT-L algorithm and visualised in Jalview^65^, with amino acid residues coloured by percentage identity. The aligned sequences included homologues from: *Serratia marcescens* (query), *Vineibacter terrae* (WP_329714985.1), *Sphingomonas* (WP_125975599.1), *Paraburkholderia sp.* (HEY4352182.1), *Paraburkholderia rhizosphaerae* (WP_134190794.1), *Caballeronia* (WP_060819512.1), *Duganella sp.* (HWW68491.1), *Pseudomonas aeruginosa* (MDX4056386.1), *Vibrio caribbeanicus* (WP_009601524.1) and *Gammaproteobacteria* (WP_023085257.1).

To visualise the relationship between conservation and structural features of Tfe2, a structural model of *S. marcescens* Tfe2 was generated using AlphaFold^66^ (version 2.3.0). Among the five output models generated, the model with the highest pLDDT score was selected for downstream analysis. This model was combined with the non-redundant multiple sequence alignment (MSA) and the conservation scores were mapped onto the 3D structure using the ConSurf server^67^. Residues were coloured by evolutionary conservation, with a magenta-green colour scheme, where magenta indicates low conservation, and green indicates high conservation. Key residues (His34, His35, Asp113 and Glu116) were highlighted as sticks in the structural representation to demonstrate that they lie within highly conserved regions, consistent with their putative catalytic or functional roles.

### Plate toxicity assay

*S. cerevisiae* WT (BH01) and Tfe2-expressing cells (KT165, JS185, JS187, JS189) were pre-grown in non-inducing liquid media (SC + 2% raffinose) at 30°C, adjusted to an OD_600_ of 1, serially diluted and 5 µl spotted onto SC + 2% raffinose agar, containing 0.02 % galactose for the induction of Tfe2 expression where indicated. Agar plates were incubated at 30°C and pictures taken after 48 h. Plate toxicity assays were repeated independently twice and representative images are shown.

### Proteomics sample preparation, LC-MS, and data processing

*S. cerevisiae* WT (BH01) and Tfe2-expressing cells (KT165) from 5 biological replicates for each strain were grown in non-inducing liquid media (SC + 2% raffinose) at 30°C to OD_600_ of 0.5 and one part was harvested for analysis of non-induced cells. The remaining cells were induced for 1 and 6 h with 1 % galactose. To account for the different growth rate of WT and Tfe2-induced cells in the presence of galactose, WT cells intended for harvesting after 6 h incubation time were set back to an OD_600_ of 0.2 before incubation with galactose. Cells were washed once in PBS, resuspended in SDS lysis buffer (50 mM TrisCl pH 8.0, 1 % SDS) and lysed by using acid-washed glass beads in a Mini bead-beater 16 (BioSpec Products, 30 sec cycles with 1 min rest time on ice, three times). Crude cell lysate was collected and SDS concentration was increased to 5 % by addition of SDS dilution buffer (50 mM TrisCl pH 8.0, 9 % SDS) at a v:v ratio of 1:1. Samples were sonicated using a UP200St ultrasonic processor (hielscher) at 90 W, 45 s pulse, 15 s rest, three times, 4 °C and clarified by centrifugation (1.000 x g, 5 min). Protein concentration was quantified using Pierce BCA Protein Assay (ThermoFisher Scientific). Samples (20 µg) were then denatured with 5 mM tris(2-carboxyethyl)phosphine (TCEP) at 60 °C for 15 minutes, alkylated with 20 mM iodoacetamide at room temperature (20 °C) for 30 minutes in the dark, and acidified to a final concentration of 2.5 % phosphoric acid. Samples were then diluted eightfold with 90 % MeOH 10 % TEAB (pH 7.2) and added to the S-trap (Protifi) micro columns. The manufacturers protocol was then followed, with a total of five washes in 90 % MeOH 10 % TEAB (pH 7.2), and trypsin added at a ratio of 1:10 enzyme:protein in 50 mM TEAB (pH 8.5) and digestion performed for 18 h at 37 °C. Peptides were dried using a vacuum concentrator and stored at −80 °C, and resuspended in 0.1 % formic acid in HPLC water immediately before mass spectrometry.

Liquid chromatography (LC) was performed using an Evosep One system with a 15 cm Aurora Elite C18 column with integrated captive spray emitter (IonOpticks), at 50 °C. Buffer A was 0.1 % formic acid in HPLC water, buffer B was 0.1 % formic acid in acetonitrile. Immediately prior to LC-MS, peptides were resuspended in buffer A and a volume equivalent to 500 ng was loaded onto the LC system-specific C18 EvoTips, according to manufacturer instructions, and subjected to the predefined Whisper-Zoom 20 SPD (where the gradient is 0-35 % buffer B, 200 nl/min, for 58 minutes, 20 samples per day were permitted) using the Evosep One (Evosep) in line with a timsToF HT mass spectrometer (Bruker) operated in diaPASEF mode. Mass and IM ranges were 300-1200 *m/z* and 0.6-1.45 1/*K_0_*, diaPASEF was performed as previously described^68^ using variable width IM-*m/z* windows without overlap. TIMS ramp and accumulation times were 100 ms, total cycle time was ∼1.8 seconds. Collision energy was applied in a linear fashion, where ion mobility = 0.6-1.6 1/K_0_, and collision energy = 20 - 59 eV.

Raw diaPASEF data files were searched using DIA-NN V 1.9^69^ using its *in silico* generated spectral library function, based on reference proteome FASTA files for *S. cerevisiae* (UP000002311, downloaded from UniProt on 24/06/2024) and a common contaminants list^70^. Trypsin specificity with a maximum of 1 missed cleavage was permitted per peptide, cysteine carbamidomethylation were set as a fixed modification. Peptide length and m/z was 7-30 and 300-1200, charge states 2-3 were included. Mass accuracy was fixed to 15 ppm for MS1 and MS2. Protein and peptide FDR were both set to 1 %. All other settings were left as defaults. Raw data and DIA-NN results files were uploaded to the proteomeXchange via the PRIDE partner repository, under PXD070762^71^. Post processing and analysis was performed using R with Rstudio. For proteomics, the protein group matrix output from DIA-NN underwent filtering to exclude contaminants, and include protein groups quantified by at least two peptides. Protein intensity was log2 transformed and normalized across samples using median normalization. For proteins present in at least 4 of 5 replicates in one condition and absent in all 5 replicates in the 2nd condition, missing values were imputed by random sampling from a downshifted normal distribution (width = 0.3, downshift = 1.8). Differential expression analysis of the log2 intensity values were subjected to processing with limma^72^, with an empirical Bayes moderated t-test.

Principle component analysis by singular value decomposition (PCA) of all strains and conditions was performed in Rstudio, using the prcomp function within the stats package to obtain principle components and the percent variance, which were ultimately plotted using the R package ggplot. Determination of ANOVA positive proteins was carried out in Perseus (v2.1.4.0) with permutation-based FDR for truncation (FDR <0.0001). Heatmap of ANOVA positive proteins was produced in R (v4.4.2) by K-means based generic clustering for 16 clusters using the R packages tidyverse (v2.0.0) and pheatmap (v.1.0.12). Line plots were created using the R package ggplot (v3.5.1). Functional enrichment analysis of Tfe2-induced cells as compared to non-induced cells was carried out in Cytoscape (version 3.10.3)^29^ applying the String network for *S. cerevisiae*. Proteins were grouped by topological clustering based on interactions of differentially regulated proteins (foldchange<2, p<0.05, MCL = 3).

### Fluorescence microscopy

For visualisation of the amino acid transporter Agp1-GFP, *S. cerevisiae* WT (JS47) and Tfe2-expressing cells (JS93), or *bul1Δ* cells expressing Tfe2 (JS399), were transformed with plasmid pAG416-P_GPD_-AGP1-yEGFP-T_CYC1_^33^ and pre-grown in non-inducing liquid media (SC-HIS-URA + 2% raffinose) at 30°C for 16 h. Cells were diluted to an OD_600_ of 0.06 (JS47, JS93) and 0.07 (JS399) in synthetic minimal medium without amino acids and ammonium sulphate, supplemented with 2% raffinose and 0.1% allantoin as sole nitrogen source and grown for 20 h at 30°C to reach mid log growth (OD_600_ = 0.5-1.0). To allow for visualisation of the vacuoles, FM4-64 was added 1 h prior to the first time point at 5 µg/ ml followed by culturing all strains and conditions in the dark. WT and Tfe2 strains were imaged before (non-induced) and after treatment with 1 % galactose for 1 and 3 h in the respective strains, with WT cells also treated, where indicated, with 10 mM alanine (Ala) or 25 µg/ml cycloheximide. To account for auxotrophies in the background strain all strains carried the pRS415-CgHIS3MET15^33^ plasmid harbouring both the CgHIS3 and MET15 genes. Cells were imaged using 1.2 % agarose pads containing 0.5 x SC + 2 % raffinose on a Zeiss Axiocam Imager.M2 with a Zeiss Plan APOCHROMAT 63x/1.40 Oil Ph3 ∞/0,17 objective. Samples were imaged using an illumination wavelength of 450-488 nm, excitation filter of 450-490 nm and emission filter of 500-550 nm for Agp1-GFP and an illumination wavelength of 505-605 nm, excitation filter of 538-562 nm and emission filter at 570-640 nm for FM4-64 using an Axiocam 503 mono camera. Widefield image data were deconvolved using Huygens (Scientific Volume Imaging) with classic MLE and automatic signal to noise ratio estimation. Images were output in 32-bit float to maintain resulting data accuracy and down sampled through linked scaling for image presentation. Finally, images were processed in ImageJ 1.54F keeping parameters constant for all images.

For visualisation of lipid droplets *S. cerevisiae* WT (BH01) and Tfe2-expressing cells (KT165) were grown in non-inducing liquid media (SC + 2% raffinose) to an OD_600_ of 0.5 at 30°C, followed by treatment with 1 % galactose for 90 min. Bodipy^TM^ 493/503 (Thermo Fisher Scientific) was added 20 min prior to each sampled timepoint at 5 µg/ ml followed by culturing all strains and conditions in the dark. Cells were imaged as specified above using excitation conditions and filter sets as for Agp1-GFP imaging without the deconvolution step.

For visualisation of the ER and nuclear structures *S. cerevisiae* JS388 and JS389 cells were grown in non-inducing liquid media (SC + 2% raffinose) to an OD_600_ of 0.6 at 30°C, followed by treatment with 1 % galactose for 3h30. Cells were imaged as specified above using excitation conditions and filter sets as for Agp1-GFP and FM4-64 imaging without the deconvolution step. All experiments were carried out three times independently and representative images are shown.

For visualisation of the mitochondrial network *S. cerevisiae* JS271 and JS287 cells were grown in non-inducing liquid media (SC + 2% raffinose) to an OD_600_ of 0.6 at 30°C, followed by treatment with 1 % galactose for 45 minutes. Yeast cells were mounted onto a 1% agarose pad (in SC media; 1% galactose) and imaged at 30°C on a Leica SP8 3X STED Microscope using an 86x (NA 1.2) water objective. Cox4-GFP was excited at 488 nm with 18% laser power and z-stacks (0.16 μm steps) were acquired through the entire volume of the cell for analysis of mitochondrial networks. Frame size was set to 1024 x 1024 and image acquisition was performed at 16-bit pixel depth and 50.05 x 50.05 nm pixel size.

Confocal z-stacks were deconvolved using Huygens (Scientific Volume Imaging) with classic MLE and automatic signal to noise ratio estimation. Images were output in 32-bit float to maintain resulting data accuracy and down sampled through linked scaling for image presentation.

Deconvolved z-stacks were converted into IMS format using the Imaris File Converter (Oxford Instruments) using standard settings. Z-stacks were then imported in Imaris v. 10.1 (Oxford Instruments) for mitochondrial volume quantification and mitochondria were segmented using the Surfaces creation tool. The quantitative data was exported for further analysis in GraphPad Prism v10.6.1.

### Amino acid sample preparation, HPLC-MS and data analysis

*S. cerevisiae* WT (BH01) and Tfe2-expressing cells (KT165) were grown in non-inducing liquid media (SC + 2% raffinose) to an OD_600_ between 0.6 and 0.8 at 30°C, followed by treatment with 1 % galactose (Tfe2) or 25 µg/ml cycloheximide (WT cells). Cells from 4 biological replicates for each strain were harvested at timepoints 0, 30 and 60 min, washed once with ice-cold PBS and flash-frozen in liquid nitrogen. Cell pellets were resuspended in 600 µl −20°C MeOH, and after addition of 600 µl ice-cold water samples were normalised to 2.5 OD_600_ equivalents suspended in 700 µl in 1:1 MeOH/H_2_O (v/v). Cells were lysed by bead beating using acid-washed glass beads in a Mini bead-beater 16 (BioSpec Products, 30 sec cycles with 1 min rest time on ice, four times), followed by sonication in a sonicator water bath for 1 min. Crude cell lysates were collected by centrifugation, cleared by further centrifugation at 8.000 rpm for 15 min at 4 °C and dried using a vacuum concentrator. Samples were reconstituted in 50 µl H_2_O of which 10 µl were added to 70 µl borate buffer (5% m/m sodium tetraborate decahydrate). After addition of 20 µl AQC solution (3 mg 6-Aminoquinolyl-N-hydroxysuccinimidyl carbamate per ml acetonitrile), the samples were briefly vortexed and incubated for 60 seconds at room temperature. Samples were then heated to 55 °C for 10 minutes, dried using a vacuum concentrator and reconstituted in H_2_O. HPLC-MS analysis was performed using an Ultimate 3000 HPLC with TripleTOF 6600 mass spectrometer (Sciex). MS Detection was via electrospray ionization (ESI) in positive ion mode using TOF MS for the quantification of each compound. Nitrogen was used as the desolvation gas and curtain gas. The following ion source conditions were used: ISVF, +4500 V; Gas1, 20 psi; Gas2 20 psi; Curtain gas, 25 psi, Temperature, 300 °C; Declustering potential, 80 V. The chromatographic separation used reversed-phase gradient chromatography on a AccQ-Tag Ultra C18 Column, 2.5 µm, 4.6 x 100 mm (Waters). The mobile phase was composed of 10mM ammonium formate and 0.5% formic acid in water (v/v) (A) and 0.5% formic acid in acetonitrile (v/v) (B). The column temperature was maintained at 45 °C and linear gradient elution was performed at 0.4 mL/min starting at 1% B, held for 0.54 min before increasing to 9.1% over 5.2 min, then to 21.2% over 2 min, then to 59.6% over 0.3 min, and finally increasing to 90% for 0.6 min, before returning to 1% B (1.3 min) for re-equilibration. The syringe wash was 50:50 water/acetonitrile (v/v). Before analysis, two injections of double blank were performed to ensure system stability and cleanliness. Data were analysed using Graphpad Prism 10 by one-way ANOVA of the respective amino acid over all strains and conditions with Tukey’s test.

### Immunoblot analysis

For determination of Sch9/Sch9-pThr^737^ levels in *S. cerevisiae,* WT (BH01) and Tfe2-expressing cells (KT165) were grown in non-inducing liquid media (SC + 2% raffinose) to an OD_600_ of 0.9 at 30°C, followed by treatment with 1 % galactose, 25 µg/ml cycloheximide or 200 ng/ml rapamycin and combinations thereof. Cells of an equivalent of OD_600_ of 9 were harvested at timepoints 0, 30 and 60 min by adding trichloroacetic acid (TCA) at a final concentration of 10 % before centrifugation and resuspension of the pellets in 10 % TCA. After 30 min incubation on ice cells were washed twice with ice-cold acetone and sonicated for 20 sec in a sonicator bath. Cell pellets were dried using a vacuum concentrator and suspended in urea-based lysis buffer (50 mM NaPO_4_ (pH7), 1 mM NaN_3_, 25 mM MOPS (pH7), 1% SDS, 3M Urea) supplemented with bromophenol blue (BPB) and 10% mercaptoethanol. Cells were lysed by bead beating using acid-washed glass beads in a Mini bead-beater 16 (BioSpec Products, 30 sec cycles with 1 min rest time on ice, three times), incubated for 10 min at 65 °C and cleared by centrifugation. Proteins were separated by 8 % SDS-PAGE and electroblotted onto nitrocellulose (Protram, Amersham Pharmacia Biotech). Primary antibodies used were polyclonal rabbit anti-Sch9 (kind gift of C. De Virgilio lab; 1:10.000) and anti-Sch9-pThr^737^ (kind gift of R. Loewith lab; 1:10.000) and peroxidase-conjugated anti-rabbit secondary antibody (Sigma-Aldrich #A6154; 1:5.000). Visualisation was performed using Pierce ECL Plus Western Blotting Substrate (ThermoFisher Scientific), the Typhoon FLA 9500 Imaging system (Amersham Pharmacia Biotech).

For determination of Sui2/Sui2-pSer^51^ levels in *S. cerevisiae,* WT (BH01) and Tfe2-expressing cells (KT165) were grown in non-inducing liquid media (SC + 2% raffinose) at 30°C, followed by treatment with 1 % galactose or 25 µg/ml cycloheximide for 30 and 60 min. Protein extracts were prepared using acid-washed glass beads by bead-beating in ice-cold lysis buffer (25 mM sodium phosphate pH 7.5, 1 mM PMSF, 1 tablet of cOmplete Mini EDTA-free protease inhibitor cocktail (Roche; 1 tablet per 10 ml of buffer)). NuPAGE™ reducing agent and NuPAGE™ LDS sample buffer were added to extracts before boiling at 95°C for 5 minutes. Proteins were separated by 4-12% Bis-Tris NuPAGE™ gels (ThermoFisher Scientific) and electroblotted onto PVDF membrane (Amersham Pharmacia Biotech). Primary antibodies used were polyclonal chicken anti-Sui2 (kind gift of G. D. Pavitt lab^73^; 1:1.000) and polyclonal rabbit anti-Sui2-pSer^51^ (Cell Signalling Technologies #9721; 1:10.000) with secondary antibodies IRDye 680RD donkey anti-chicken and IRDye 800CW goat anti-rabbit IgG (LI-COR Biosciences #926-68072 and #926-32211; 1:10.000 each). Visualisation was performed using the LI-COR ODYSSEY CLx Imaging system (LICORbio). Fluorescence signals were quantified using 3 independent imaging blots and the Image Studio software.

### Polysome profiling and analysis

Polysome profiling was performed as previously described^74^. Briefly, *S. cerevisiae* WT (BH01) and Tfe2-expressing cells (KT165) were grown in non-inducing liquid media (SC + 2% raffinose) to an OD_600_ of 0.6 to 0.8 at 30°C, followed by treatment with 1 % galactose or 25 µg/ml cycloheximide for 30 and 60 minutes. Cells were transferred to Falcon tubes containing cycloheximide (final conc. 0.1 mg/ml) and rapidly cooled on ice for 10 minutes before centrifugation. Cell pellets were washed and lysed by acid-washed glass beads in lysis buffer (20 mM HEPES pH 7.4, 100 mM potassium acetate, 2 mM magnesium acetate, 0.5 mM DTT, 0.4 mM cycloheximide; 20 sec cycles with 40 seconds rest time on ice, eight times). Supernatant was collected by centrifugation and 2.5 *A*_260_ units were layered onto 15-50% sucrose gradients. Gradients were separated by ultracentrifugation in an SW41 Ti rotor (2.5 h at 278 000 × *g*, 4°C). Polysome profiles were generated using TRIAX™ flow cell UV/Vis recorder continuously monitoring *A*_260_. Polysome traces and polysome to monosome ratios were generated using the QuAPPro (v. 0.3.1) online tool.

### Metabolic labelling with NBD-C_6_-ceramide and analysis by thin layer chromatography (TLC)

*S. cerevisiae* WT (BH01) and Tfe2-expressing cells (KT165) were grown in non-inducing liquid media (SC + 2% raffinose) to an OD_600_ of 0.6 to 0.8 at 30°C, followed by treatment with 1 % galactose or 0.25 µg/ml AureobasidinA as control condition for 2 hours. Cells were normalized to 2.5 OD_600_ equivalents/ml, supplemented with 5 μM of NBD C_6_-ceramide conjugated to bovine serum albumin (Thermo Fisher Scientific #11519126) and incubated for 2 h at 30 °C. Yeast were harvested by centrifugation and washed twice with phosphate buffered saline. 400 µl of chloroform/methanol (1:1, v/v) was added and cells were lysed by using acid-washed glass beads (30 sec cycles with 1 min rest time on ice, four times). The lysate was centrifuged, the supernatant transferred to a new reagent tube and the pellet re-extracted with 400 µl chloroform/methanol/water (10:10:3). Samples were centrifuged and the supernatant pooled with the previous extract. The lipid fraction was isolated by phase separation by addition of H_2_O and dried in a vacuum concentrator. TLC analysis was performed as previously described^75^. Briefly, reaction products were re-suspended in 20 μl of chloroform/methanol/water (10:10:3) and cell mass equivalents fractionated using HPTLC silica gel 60 plates (Merck) and the eluent system chloroform:methanol:aqueous 0.25% KCl (55:45:10). Imaging and quantification was carried out using the Typhoon FLA 9400 Imaging system (Amersham Pharmacia Biotech).

### Lipid extraction and lipidomic analysis by LC-MS

*S. cerevisiae* WT (BH01) and Tfe2-expressing cells (KT165), 6 biological replicates per strain, were grown in non-inducing liquid media (SC + 2% raffinose) to an OD_600_ between 0.4 and 0.6 at 30°C, followed by treatment with 1 % galactose for 6 hours. The equivalent of 25 OD_600_ units of cells were harvested by centrifugation, washed twice in ammonium carbonate buffer (150 mM, pH 8.0) and frozen in liquid nitrogen. Cell pellets were suspended in ammonium carbonate buffer and normalised to 20 OD_600_ units/ml. Cells were lysed using acid-washed glass beads by vortexing (1 min cycles with 1 min rest time on ice, 10 times) and the lysate was collected by centrifugation. Samples were vortexed before transferring 100 µl of lysate to 1 ml of MeOH. Lipids were extracted as described previously^76^. Briefly, samples were incubated with 6 ml of chloroform/methanol (2/1, v/v) for 1 hour at 4 °C. Samples were partitioned by the addition of 0.1 M KCl and the mixture was centrifuged to facilitate phase separation. The lower chloroform layer was evaporated to dryness under nitrogen gas and reconstituted in methanol containing 5 mM ammonium formate.

Lipids were analysed by liquid chromatography-mass spectrometry (LC-MS) using a Thermo Exactive Orbitrap mass spectrometer equipped with a heated electrospray ionization (HESI) probe and interfaced with a Dionex UltiMate 3000 RSLC system (Thermo Fisher Scientific, Hemel Hempstead, UK). Samples (10 µL) were injected onto a Thermo Hypersil Gold C18 column (2.1 mm x 100 mm; 1.9 μm) maintained at 50 °C. Mobile phase A consisted of water containing 10 mM ammonium formate and 0.1% (v/v) formic acid. Mobile phase B consisted of a 90:10 (v:v) mixture of isopropanol:acetonitrile containing 10 mM ammonium formate and 0.1% (v/v) formic acid. The initial conditions for analysis were 65% mobile phase A, 35% mobile phase B and the percentage of mobile phase B was increased from 35 to 65% over 4 min, followed by 65% to 100% over 15 min, with a hold for 2 min before re-equilibration to the starting conditions over 6 min. The flow rate was 400 μL/min. All samples were analysed in positive and negative ionisation modes over the mass-to-charge ratio (m/z) range of 250 to 2,000 at a resolution of 100,000. Raw LC-MS data was processed with Progenesis QI v2.4 software (Non-linear Dynamics). Signals were normalised using all ion normalisation, before retention and mass aligned features were annotated using the LipidMaps database with a mass error tolerance of 5 ppm. Initial searches were performed for the molecular ions of CL [M+H]^+^, Cers [M+Na]^+^, DG [M+Na]^+^, IPC [M-H]^-^, LPC [M+H]^+^, LPE [M+H]^+^, MIPC [M-H]^-^, PC [M+H]^+^, PE [M+H]^+^, PI [M-H]^-^ and TG [M+Na]^+^. Annoations were then confirmed using additional species specific ESI adduct formations and RPLC retention characteristics. Data were analysed using Graphpad Prism 10 by multiple unpaired t-tests with Holm-Šídák’s test.

### Lipid-targeting antifungal drug susceptibility assay and FACS analysis

*S. cerevisiae* WT (BH01) and Tfe2-expressing cells (KT165) were grown in non-inducing liquid media (SC + 2% raffinose) to an OD_600_ of 0.4 at 30°C, induced with 1 % galactose for 5h30, followed by treatment with 500 ng/ml AureubasidinA, 40 µg/ml Myriocin or 2.5 µg/ml Miltefosine for 16 hours. DMSO was added to galactose-incubated cells as control condition. Cells were washed once in SC media and normalised to an OD_600_ of 0.3, of which 200 µl were transferred into a round-bottomed 96-well plate. Cells were stained with propidium iodide (PI; 10 µM final concentration) for 15 min at room temperature in the dark. Samples were analysed without washing. Unstained and untreated controls were prepared for both wild-type and Tfe2-expressing strains, and vehicle controls containing DMSO only were included. As a PI-positive control an additional Aureobasidin A treated wildtype sample was used. Flow cytometry was performed on a BD FACSymphony™ A3 Cell Analyzer (BD Biosciences) equipped with a high-throughput sampler, and data collected through the BD FACSDiva (v9.0) software. PI fluorescence was excited using the 488 nm and 561 nm lasers, and emission was collected using a 610/20 nm bandpass filter. Isolation of singlet yeast cells, and exclusion of debris, was gated for using FSC and SSC parameters. Fluorescence gates were established using the positive and negative controls to define background auto-fluorescence, and PI-positive cell boundaries. For each sample, 50,000 events were recorded. Data analysis was carried out using FCS Express 7 (Research Edition). The percentage of PI-positive cells was calculated against total events present within population gates for each biological replicate, and averaged across three independent replicates collected on separate days.

### EM structural analysis

*S. cerevisiae* WT (BH01) and Tfe2-expressing cells (KT165) were grown in non-inducing liquid media (SC + 2% raffinose) to an OD_600_ of 0.6 at 30°C and treated with 1 % galactose for 2h30. For conventional TEM analysis cell cultures were chemically fixed with 2% glutaraldehyde and 2% paraformaldehyde in PHEM buffer (60 mM PIPES, 25 mM HEPES, 10 mM EGTA, 2 mM MgCl_2_, pH 7.3) for 1 h, and kept at 4 °C until further processing. Samples were washed 3 times in Phem buffer for 5 minutes, incubated for 1 h in 1% Na-metaperiodate in PBS, and washed 3 times in PBS for 5 minutes. For post-fixation, cells were incubated for 1 h in 2% potassium permanganate in dH_2_O, followed by 3 washed in dH_2_O for 5 min. Samples were subsequently passed through a gradient of dimethylformamide (DMF; 30%, 50%, 70%, 80%, 90%) for 10 minutes each, followed by 2 incubations in 100% DMF, 2 times for 15 min each, and embedded gradually in SPURR resin over 2 days. 70 nm ultrathin sections were collected on formvar-coated copper EM grids, contrasted in Reynold’s lead citrate and imaged using a JEOL JEM 1400 transmission electron microscope operated at 120 kV and equipped with a Rio16 4k x 4k digital camera (Gatan, Ametek, Leicester, UK).

For cryo-fixation and vitrification of samples, cell cultures were passed through a 0.2 um polycarbonate filter by applying a mild vacuum and concentrated cells were loaded into high pressure freezer (HPF) aluminium carriers. Samples were high pressure frozen in a HPM Live µ (CryoCapCell, Aubière, France) and freeze substituted with 1% osmium tetroxide in acetone using a AFSM2 freeze substitution unit (Leica Microsystems, Milton Keynes, UK). Samples were then washed in pure acetone and embedded over 2 days in SPURR resin. Sectioning and imaging were done as described above.

To quantify the density of ribosomes in the cytosol, high pressure frozen cells were imaged at random at a nominal magnification of 20k X. Ribosome density was estimated by spatial stereology utilising the point counting method ^77^. A square lattice grid was placed over the images at random in ImageJ and points counted over ribosome density as well as the cytosol (grid spacing 0.1 um for ribosomes, 0.2 um for cytosol) to estimate the fractional density of ribosomes per cytosol from the sampled images.

To measure the width of inner and outer cell wall, high pressure frozen cells were imaged at random at a nominal magnification of 20k X. Cell wall width was measured for a minimum of 43 images per strain using the line tool in ImageJ. For each cell image, 10 measurements were randomly taken along the inner and outer wall.

### Monosaccharide quantification using HPIC

Cell wall composition was analysed as described previously^78^. Briefly, *S. cerevisiae* WT (BH01) and Tfe2-expressing cells (KT165) were grown in non-inducing liquid media (SC + 2% raffinose) to an OD_600_ of 0.6 at 30°C and treated with 1 % galactose for 2h30. Cells were harvested by centrifugation, washed twice in water and lysed using a FastPrep machine (MP Biomedicals). The homogenate was pelleted and washed with 1M NaCl to remove proteins and heated for 10 mins at 100 °C in SDS extraction buffer (500mM Tris-HCL, pH7.5, 2% (wt/vol) SDS, 0.3 M β-mercaptoethanol, 1 mM EDTA), before freeze drying. β-glucan, mannan and chitin levels were determined by hydrolysis and quantification of glucose, mannose and glucosamine, respectively, using 2M Trifluoroacetic acid at 100 °C for 3 h. The hydrolysates were analysed by High-Pressure-Ion-Chromatography as described previously^79^, with the following modifications. 0.4 μl of sample was injected into a Dionex carbohydrate analyser equipped with a CarboPac PA20 column (0.4×150mm), guard column and an ED50 Pulsed amperometric detector (PAD). The samples were eluted with a gradient of 5-100 mM at a flow rate of 0.08 ml/min for 25 minutes.

## Acknowledgements

This work was supported by the Wellcome Trust (215599/Z/19/Z, J.Q., S.J.C, M.T., N.A.R.G., & C.R.). We thank Professor Hiroshi Takagi (Nara Institute of Science and Technology, Japan) for the kind gift of plasmids to allow Agp1-GFP localisation, and Professor Claudio De Virgilio (University of Fribourg, Switzerland) for the kind gift of anti-Sch9 and anti-Sch9-pThr^737^ antibodies

## Author Contributions

K.T., J.Q., S.J.C. and G.P. conceived the study. K.T., G.P., A.A., V.S., A.F., P.P., J.H., I.L., M.L., F.L. and C.H., performed experimental work and analysed data. K.T., G.P., P.W.D., C.H., P.D.W., N.A.R.G., C.R., M.T., C.M.G., S.J.C. and J.Q., designed experiments, analysed data and interpreted data. K.T., G.P. and J.Q. wrote the manuscript with contributions and input from the other authors.

