## Supplementary Table 5 for "The Type VI Secretion System Antifungal Effector Tfe2 Inhibits Protein Translation and Drives Hyperactivation of TORC1"

Supplementary Table 5. Strains and Plasmids used in this study.

| **Name** | **Description** | **Source/**  **Reference** |
| --- | --- | --- |
| **Strains** |  |  |
| *S. cerevisiae* | |  |
| K699 | *MAT***a** *ade2-1* *trp1-1 leu2-3*,*112 his3-11*,*15 ura3 can1-100* | ^1^ |
| BH01 | K699 *his3*::P*_GAL1_* | ^2^ |
| KT165 | K699 *his3*::P*_GAL1_*-*SMDB11*_*1083* | ^2^ |
| JS185 | K699 *his3*::P*_GAL1_*-*SMDB11*_*1083*(H34A) | This study |
| JS187 | K699 *his3*::P*_GAL1_*-*SMDB11*_*1083*(D113A) | This study |
| JS189 | K699 *his3*::P*_GAL1_*-*SMDB11*_*1083*(H34AD113A) | This study |
| JS213 | K699 *his3*::P*_GAL1_*-*SMDB11*_*1083*-*EGFP* | This study |
| JS215 | K699 *his3*::P*_GAL1_*-*SMDB11*_*1083*(H34A)-*EGFP* | This study |
| JS219 | K699 *his3*::P*_GAL1_*-*SMDB11*_*1083*(D113A)-*EGFP* | This study |
| JS221 | K699 *his3*::P*_GAL1_*-*SMDB11*_*1083*(H34AD113A)-*EGFP* | This study |
| JS268 | K699 *his3*::P*_GAL1_* P*_EMC1_*::*EMC1*-2*GFP* | This study |
| JS284 | K699 *his3*::P*_GAL1_*-*SMDB11*_*1083* P*_EMC1_*::*EMC1*-2*GFP* | This study |
| JS388 | K699 *his3*::P*_GAL1_* P*_EMC1_*::*EMC1*-2*GFP* P*_NAB2_*::*NAB2*-*mCherry* | This study |
| JS389 | K699 *his3*::P*_GAL1_*-*SMDB11*_*1083* P*_EMC1_*::*EMC1*-2*GFP* P*_NAB2_*::*NAB2*-*mCherry* | This study |
| JS271 | K699 *his3*::P*_GAL1_* P*_COX4_*::*COX4*-*GFP* | This study |
| JS287 | K699 *his3*::P*_GAL1_*-*SMDB11*_*1083* P*_COX4_*::*COX4*-*GFP* | This study |
| BY4741 | *MAT***a** *his3*Δ*1* *leu2*Δ*0* *met15*Δ*0* *ura3*Δ*0* | ^3^ |
| JS47 | BY4741 *his3*::P*_GAL1_* | This study |
| JS93 | BY4741 *his3*::P*_GAL1_*-*SMDB11*_*1083* | This study |
| JS393 | BY4741 *bul1*Δ::KanMX6 | ^4^ |
| JS398 | BY4741 *bul1*Δ::KanMX6 *his3*::P*_GAL1_* | This study |
| JS399 | BY4741 *bul1*Δ::KanMX6 *his3*::P*_GAL1_*-*SMDB11*_*1083* | This study |
| *S. marcescens* | |  |
| YL37 | Db10 Δ9 [Δ*ssp1* (Δ*SMDB11_2261*), Δ*ssp2* (Δ*SMDB11_2264*), Δ*ssp3/tfe1* (Δ*SMDB11_1112*), Δ*ssp4* (Δ*SMDB11_3980*), Δ*ssp5* (Δ*SMDB11_4628),* Δ*ssp6* (Δ*SMDB11_4673*), Δ*rhs1* (Δ*SMDB11_2278*), *rhs2_H1369A_* (*SMDB11_1610_H1369A_*), Δ*slp* (Δ*SMDB11_0927*)] | ^5^ |
| YL39 | Db10 Δ9 Δ*tfe2* (Δ*SMDB11_1083*) | This study |
| **Plasmids** |  |  |
| pSC1384 | pGED1 derived plasmid carrying empty PGAL1 promoter construct (integrative) | ^2^ |
| pSC1380 | pGED1 derived plasmid for galactose-inducible expression (PGAL1) of SMDB11_1083 (TFE2) (integrative) | ^2^ |
| JE414 | pGED1 derived plasmid for galactose-inducible expression (PGAL1) of SMDB11_1083(H34A) (integrative) | This study |
| JE415 | pGED1 derived plasmid for galactose-inducible expression (PGAL1) of SMDB11_1083(D113A) (integrative) | This study |
| JE416 | pGED1 derived plasmid for galactose-inducible expression (PGAL1) of SMDB11_1083(H34AD113A) (integrative) | This study |
| JE424 | pGED1 derived plasmid for galactose-inducible expression (PGAL1) of SMDB11_1083-EGFP (integrative) | This study |
| JE425 | pGED1 derived plasmid for galactose-inducible expression (PGAL1) of SMDB11_1083(H34A)-EGFP (integrative) | This study |
| JE427 | pGED1 derived plasmid for galactose-inducible expression (PGAL1) of SMDB11_1083(D113A)-EGFP (integrative) | This study |
| JE428 | pGED1 derived plasmid for galactose-inducible expression (PGAL1) of SMDB11_1083(H34AD113A)-EGFP (integrative) | This study |
| pAG416-P_GPD_-AGP1-yEGFP-T_CYC1_ | pRS416-ccdB-yEGFP-derived plasmid for expression of Agp1-yEGFP | ^6^ |
| pRS415-CgHIS3MET15 | pRS415-derived plasmid for expression of His3 and Met3 | ^6^ |
| ZJOM22 | p1k-EMC1-2GFP; plasmid for expression of Emc1-2GFP under the native promoter (integrative) | ^7^ |
| ZJOM82 | p1k-NAB2-mCherry; plasmid for expression of Nab2-mCherry under the native promoter (integrative) | ^7^ |
| ZJOM151 | p1k-COX4-GFP; plasmid for expression of Cox4-GFP under the native promoter (integrative) | ^7^ |
