## Supplementary Figure 1 for "The Type VI Secretion System Antifungal Effector Tfe2 Inhibits Protein Translation and Drives Hyperactivation of TORC1"

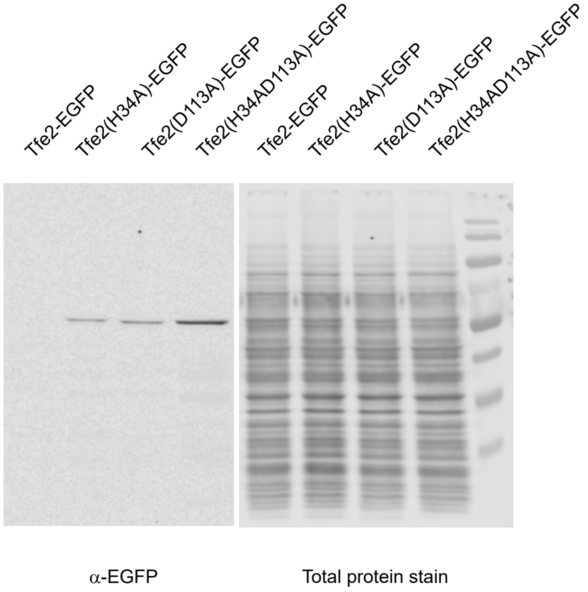


**Supplementary Figure 1.** Immunoblot of the WT Tfe2-EGFP, the single amino acid variants Tfe2(H34A)-GFP and Tfe2(D113A)-GFP and the double mutant variant Tfe2(H34AD113A)-GFP after induction with 0.2 % galactose for 1 h. Lysates were analysed using anti-GFP (α-EGFP) antibody and Revert 700 total protein stain (Licor #926-11011).
