## Supplementary Figure 2 for "The Type VI Secretion System Antifungal Effector Tfe2 Inhibits Protein Translation and Drives Hyperactivation of TORC1"

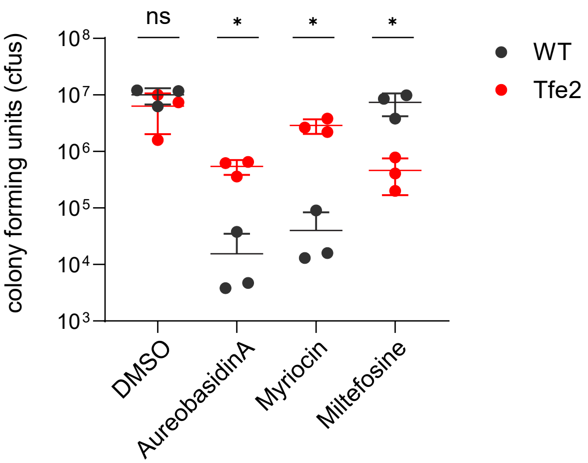


**Supplementary Figure 2.** Impact of Tfe2 toxication on susceptibility to lipid-targeting antifungal drugs. Control and Tfe2-cells were grown in the presence of 1 % galactose for 5h30 and treated with 500 ng/ml AureubasidinA, 40 µg/ml Myriocin and 2.5 µg/ml Miltefosine with DMSO as control for 16 hours. Cell survival was calculated by determination of colony forming units. Data are presented as mean ± SD with individual data points overlaid (n=3 biological replicates; * P<0.05; ns not significant; multiple unpaired t-tests with Holm-Šídák's test).
