## Supplementary Figure 3 for "The Type VI Secretion System Antifungal Effector Tfe2 Inhibits Protein Translation and Drives Hyperactivation of TORC1"

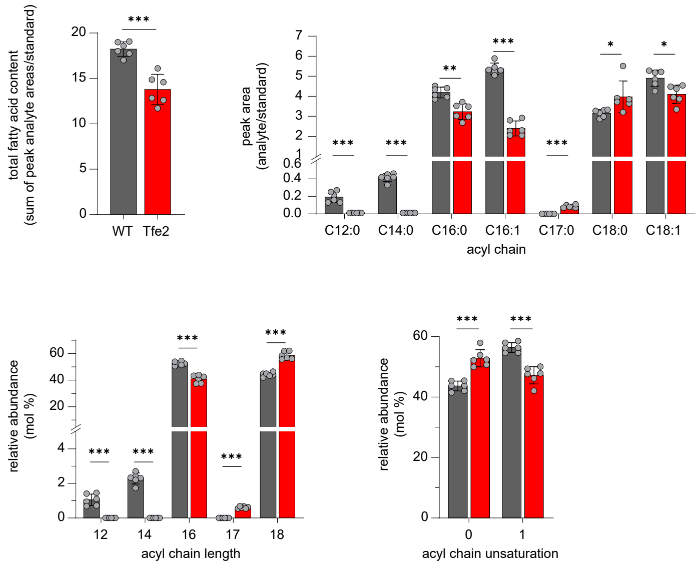


**Supplementary Figure 3.** Changes in acyl chain composition to Tfe2-intoxication as determined by GC-MS/MS analysis, shown as total fatty acid content, single acyl chain species, combined acyl chain length species and unsaturation levels. Data are presented as mean ± SD with individual data points overlaid (n=6 biological replicates; *** P<0.001, ** P<0.01,* P<0.05; ns not significant; multiple unpaired t-tests with Holm-Šídák’s test).
