## Supplementary Figure 4 for "The Type VI Secretion System Antifungal Effector Tfe2 Inhibits Protein Translation and Drives Hyperactivation of TORC1"

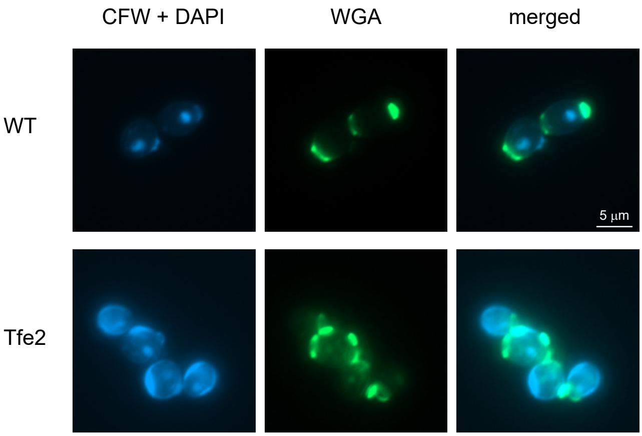


**Supplementary Figure 4.** Fluorescence microscopy of lateral cell wall (blue) and bud scar situated chitin (green) in yeast control (WT) and Tfe2-inducing strains by Calcofluor White (CFW; blue) and wheat germ agglutinin-FITC (WGA; green) staining. Cells were grown in the presence of 1 % galactose for 4 h.
