## Supplementary Figure 5 for "The Type VI Secretion System Antifungal Effector Tfe2 Inhibits Protein Translation and Drives Hyperactivation of TORC1"

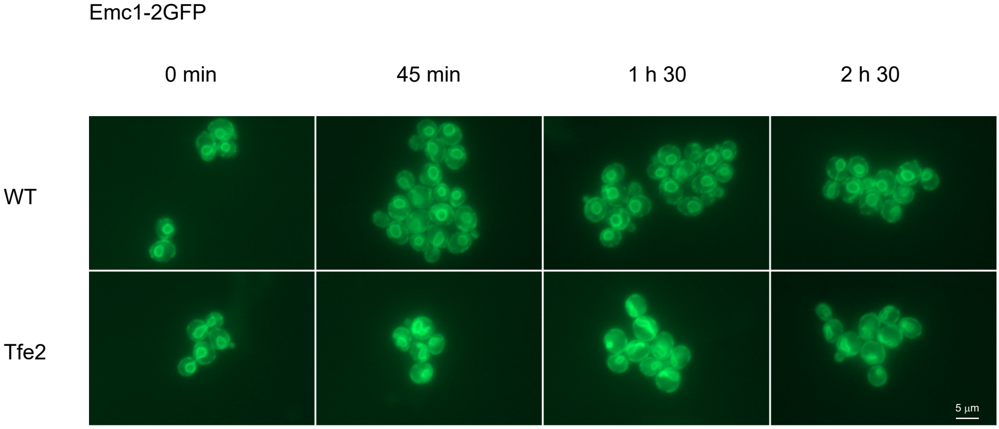


**Supplementary Figure 5.** Fluorescence microscopy of ER structures in yeast control (WT) and Tfe2-inducing strains carrying the chromosomally integrated organelle-specific fluorophore-tagged protein Emc1-2GFP. Cells were grown in the presence of 1 % galactose for the timepoints indicated.
