## Supplementary Figure 6 for "The Type VI Secretion System Antifungal Effector Tfe2 Inhibits Protein Translation and Drives Hyperactivation of TORC1"

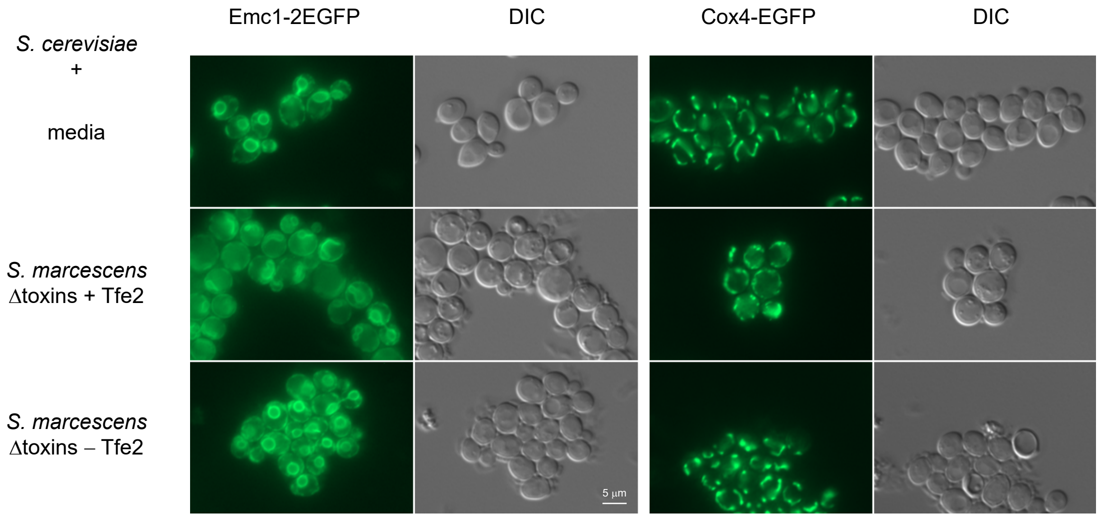


**Supplementary Figure 6.** Fluorescence microscopy of ER and mitochondrial structures in *S. cerevisiae* cells carrying the chromosomally integrated organelle-specific fluorophore-tagged proteins Emc1-2EGFP (ER) and Cox4-EGFP (mitochondria). Cells were visualised after 7h30 co-culture with media only or *S. marcescens* deleted in all known T6SS effector proteins (Δtoxins) except Tfe2 (+Tfe2) or additional deletion of Tfe2 (-Tfe2) .
